# Ectopic engraftment of nociceptive neurons derived from hPSCs for pain relief and joint homeostasis

**DOI:** 10.64898/2025.12.16.694733

**Authors:** Zhuolun Wang, Weixin Zhang, Ju Wang, Zhiping Wu, Xu Cao, Junmin Peng, Gabsang Lee, Xinzhong Dong

## Abstract

Chronic pain arises from the interplay of inflammatory signals that activate and sensitize nociceptors within injured tissues. Most analgesics fail clinically due to their mono-targeted mechanisms. Here, we apply human pluripotent stem cell–derived nociceptive neurons (hPSC-NNs) as therapeutic agents for osteoarthritis, targeting both pain and joint degeneration. We generated sensory neurons from hPSCs and identified *CD200* as a nociceptor marker. Transcriptomic and functional profiling revealed that *CD200^high^*hPSC-NNs closely resemble human nociceptors, expressing pain-relevant receptors and ion channels. Strikingly, ectopic transplantation of *CD200^high^*hPSC-NNs into the knee joint of osteoarthritic mice reduced pain and promoted bone and cartilage repair, whereas *CD200^low^* cells exhibited no benefit. Mechanistically, human and mouse proteomics suggest that *CD200^high^*hPSC-NNs act as decoys by sequestering inflammatory ligands while secreting reparative factors in joint tissues. These findings uncover a fundamental role of nociceptors in tissue repair, providing a multi-targeted, disease-modifying strategy for OA and chronic pain.

**GRAPHIC ABSTRACT:** 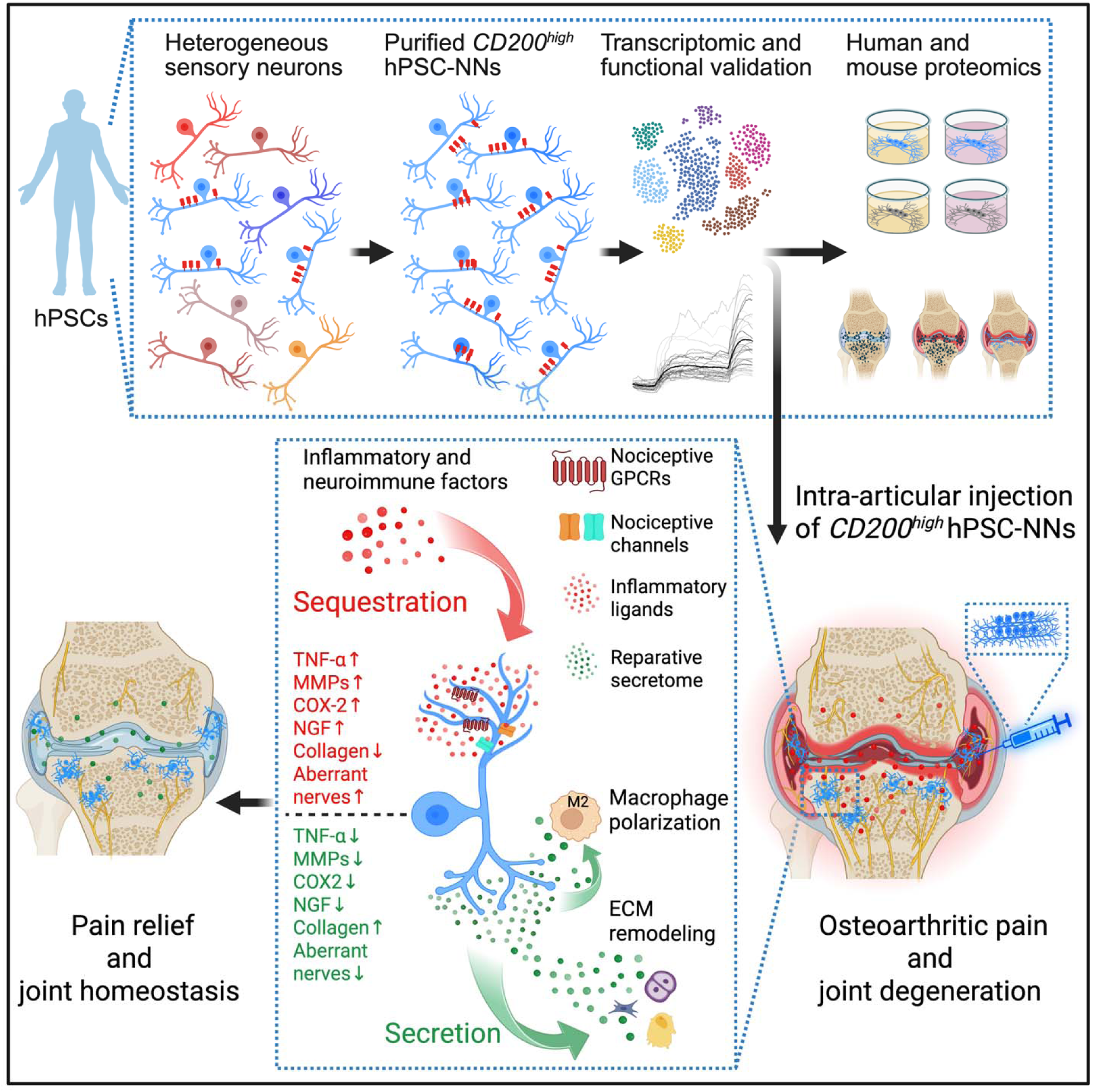

**HIGHLIGHTS:**

- hPSC-derived nociceptors (hPSC-NNs) as Decoy Engraftment for Cellular Interception and Repair (DECIR) when transplanted into the knee joint, extending beyond conventional regenerative strategies
- *CD200* serves as a clinically actionable surface marker for the purification of hPSC-NNs
- Ectopic grafting of *CD200^high^* hPSC-NNs delivers dual benefits, alleviating pain and modulating the neuro-immune environment within joint tissues
- Proteomic analyses reveal that *CD200^high^*hPSC-NNs sequester inflammatory mediators and secrete reparative factors to support joint homeostasis

## INTRODUCTION

The human somatosensory system is comprised of diverse subtypes of sensory neurons, with each specialized in detecting specific modalities such as touch, pressure, temperature, vibration, pain, and body position. Among these, nociceptors detect potentially harmful stimuli, including noxious chemical, thermal, and mechanical stimuli, through their heterogeneous expression of ion channels and receptors. In pathological states, however, persistent exposure to pro-inflammatory mediators, including cytokines such as TNF-α, PGE2, and IL-6 and neurotrophic factors like NGF, induces nociceptor hyperexcitability and upregulation of pro-nociceptive genes, which drive peripheral and central sensitization and result in chronic pain that is refractory to traditional mono-targeted analgesics^1^.

Osteoarthritis (OA) exemplifies the convergence of inflammation and chronic pain. In early osteoarthritis, inflammatory cytokines drive osteoclast-mediated subchondral bone loss, while later stages are marked by abnormal osteoblast activity and development of subchondral sclerosis. This altered microenvironment creates a self-perpetuating axis that promotes aberrant nerve sprouting, amplifies pain, and sustains joint inflammation^2^. Although the inflammatory nature of OA-associated pain is well recognized, current therapeutic strategies, including nonsteroidal anti-inflammatory drugs (NSAIDs), COX2 inhibitors, and anti-NGF antibodies^3,4^, target singular pathways and fail to address the complex neuro-immune interactions that drive disease progression.

Human pluripotent stem cell (hPSC)-derived neurons hold significant promise in regenerative medicine, primarily through replacement of damaged neurons in neurodegenerative diseases^5^. Traditional regenerative approaches involve transplanting neurons into their native niches, facilitating replacement of lost or injured cells. However, this paradigm does not address conditions involving complex tissue environments driven by neuro-immune interactions, such as OA.

Here, we introduce an alternative therapeutic strategy — Decoy Engraftment for Cellular Interception and Repair (DECIR) — by ectopically grafting purified hPSC-derived nociceptive neurons (hPSC-NNs) directly into the knee joints of mice. Distinct from conventional regenerative therapies, these transplanted hPSC-NNs survive and function within a non-native environment, actively modulating the pathological neuro-immune axis underlying OA. Given their intrinsic responsiveness to inflammatory stimuli – due to robust expression of receptors and ion channels characteristic of human nociceptors – hPSC-NNs can serve as biological decoys. We hypothesize that DECIR can locally sequester inflammatory mediators and reshape the joint microenvironment, providing dual therapeutic benefits: alleviation of chronic pain and reversal of OA-associated joint pathology.

## RESULTS

### Isolation and Functional Characterization of Molecularly Defined Subsets of Nociceptive and Pruriceptive hPSC-SNs

We generated hPSC-SNs from hPSCs using an optimized differentiation protocol. First, dua⍰lSMAD inhibition via concurrent blockade of BMP and TGF⍰β signaling was used to direct hPSCs toward a neural crest fate^6^. Next, we exposed the neural crest progenitors to a cocktail of neurotrophic factors (e.g., NGF and GDNF) to bias differentiation toward nociceptor⍰like sensory neurons (Figures 1A and 1B). To isolate molecularly defined subsets of hPSC-SNs, we generated *TRPV1::GFP*, *SCN9A::GFP*, and *MRGPRX1::GFP* reporter lines (Figure 1C). TRPV1 and SCN9A label small- and medium-diameter nociceptors, while MRGPRX1 marks a non-peptidergic subset specialized in pruritogen detection^7,8^. The fidelity of each reporter line was validated by fluorescence-activated cell sorting (FACS) for GFP expression at 70 days *in vitro* (DIV; Figure 1D) and further confirmed by bulk RNA sequencing (Figure 2B). The spatial distribution of *TRPV1*+, *SCN9A*+, and *MRGPRX1*+ hPSC-SNs was visualized using anti-GFP staining (Figure 1E). To determine the temporal dynamics of subtype specification, we performed longitudinal FACS analyses across all three reporter lines. *MRGPRX1*+ neurons were not robustly detected until around 35 DIV, whereas *TRPV1*+ and *SCN9A*+ populations did not exhibit strong expression until after 70 DIV (Figure 1F). In summary, we generated and validated purifiable subsets of *TRPV1::GFP*, *SCN9A::GFP*, and *MRGPRX1::GFP* reporter hPSC-SNs.

**Figure 1.**
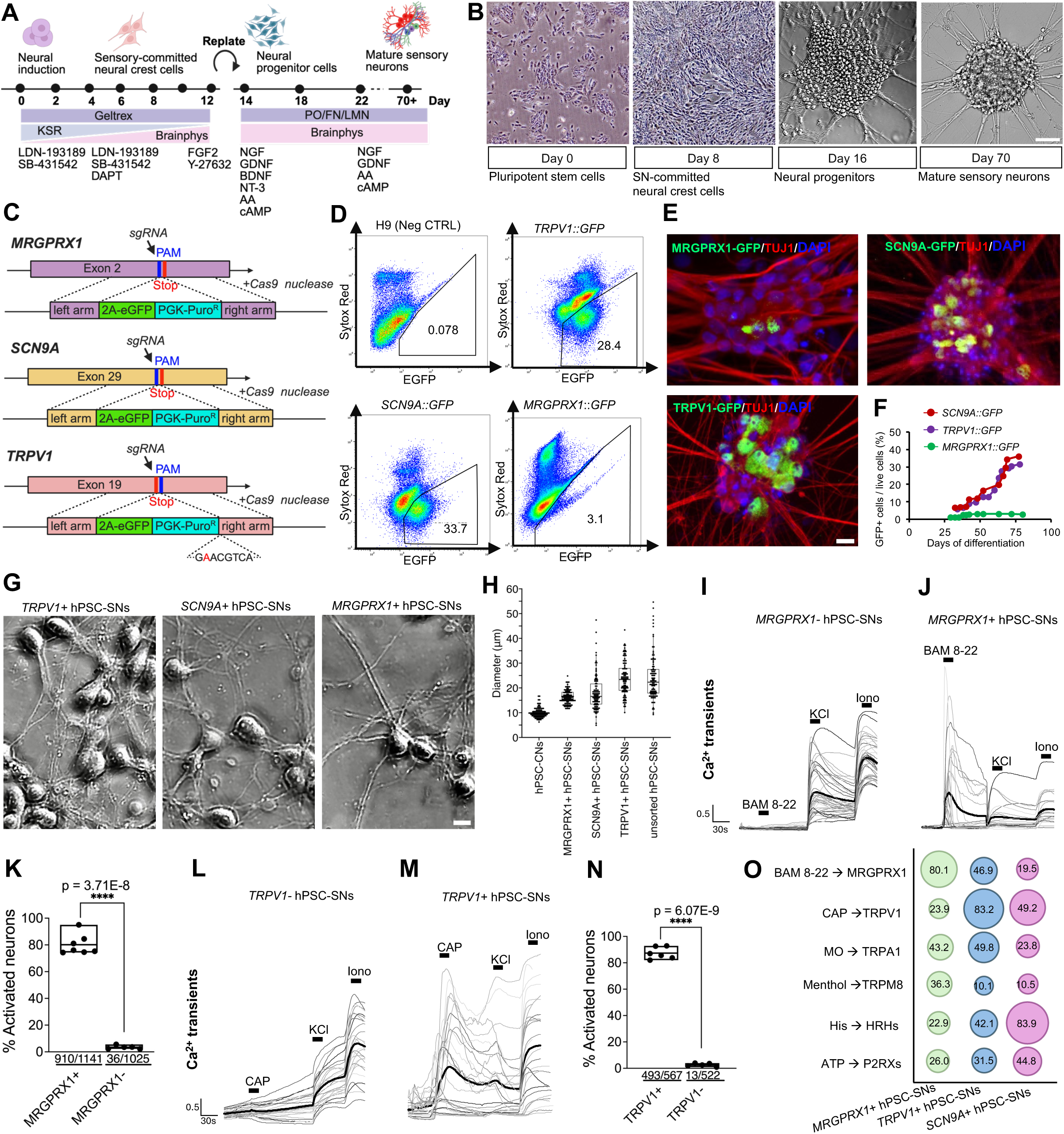
Isolation and functional characterization of *MRGPRX1::GFP, SCN9A::GFP*, and *TRPV1::GFP* hPSC-SNs. **(A)** Optimized protocol for differentiation of hPSCs into hPSC-SNs. **(B)** Bright-field images at key stages of differentiation; scale bar, 100μm. **(C)** Schematic of CRISPR–Cas9 knock-in constructs generating *MRGPRX1::GFP, SCN9A::GFP*, and *TRPV1::GFP* reporter lines. **(D)** FACS analysis of GFP expression in hPSC-SNs from each reporter line and unedited H9 controls (n = 3 clones). **(E)** Proportion of GFP+ neurons in *MRGPRX1::GFP, SCN9A::GFP*, and *TRPV1::GFP* unsorted cultures, quantified by co-immunostaining for eGFP (green) and TUJ1 (red); scale bar, 10 μm. **(F)** Time course of GFP+ cell expression in *MRGPRX1::GFP, SCN9A::GFP*, and *TRPV1::GFP* hPSC-SNs measured by FACS (n = 5 independent differentiations). **(G)** Representative images of FACS-purified *TRPV1+, SCN9A*+, and *MRGPRX1+* hPSC-SNs; scale bar, 10μm. **(H)** Boxplot of soma diameters measured in hPSC-derived cortical neurons (hPSC-CNs), FACS-purified *MRGPRX1+*, *SCN9A*+, *TRPV1+* hPSC-SNs and unsorted hPSC-SNs (n = 100 cells per group); boxes indicate the interquartile range (25th–75th percentile) and horizontal lines denote medians. **(I-K)** Calcium transients (F_t_/F_₀_) in *MRGPRX1-* **(I)** versus *MRGPRX1*+ **(J)** neurons in response to 10 μM BAM8-22; bold traces show the mean, with **(K)** quantifying the proportion of activated cells among all KCl-responsive neurons (n ≥ 5 independent differentiations; unpaired two-tailed t-test; mean ± range with individual values overlaid). **(L-N)** Calcium transients in *TRPV1*-**(L)** versus *TRPV1*+ **(M)** neurons upon 10 μM capsaicin; bold traces show the mean, with **(N)** quantifying the proportion of activated cells among all KCl-responsive neurons (n ≥ 5 independent differentiations; unpaired two-tailed t-test; mean ± range with individual values overlaid). **(O)** Percentage of FACS-purified hPSC-SNs responding to BAM8-22 (10μM), capsaicin (10μM), mustard oil (100μM), menthol (100μM), histamine (50μM), and ATP (50μM) (n ≥ 5 independent differentiations).

**Figure 2.**
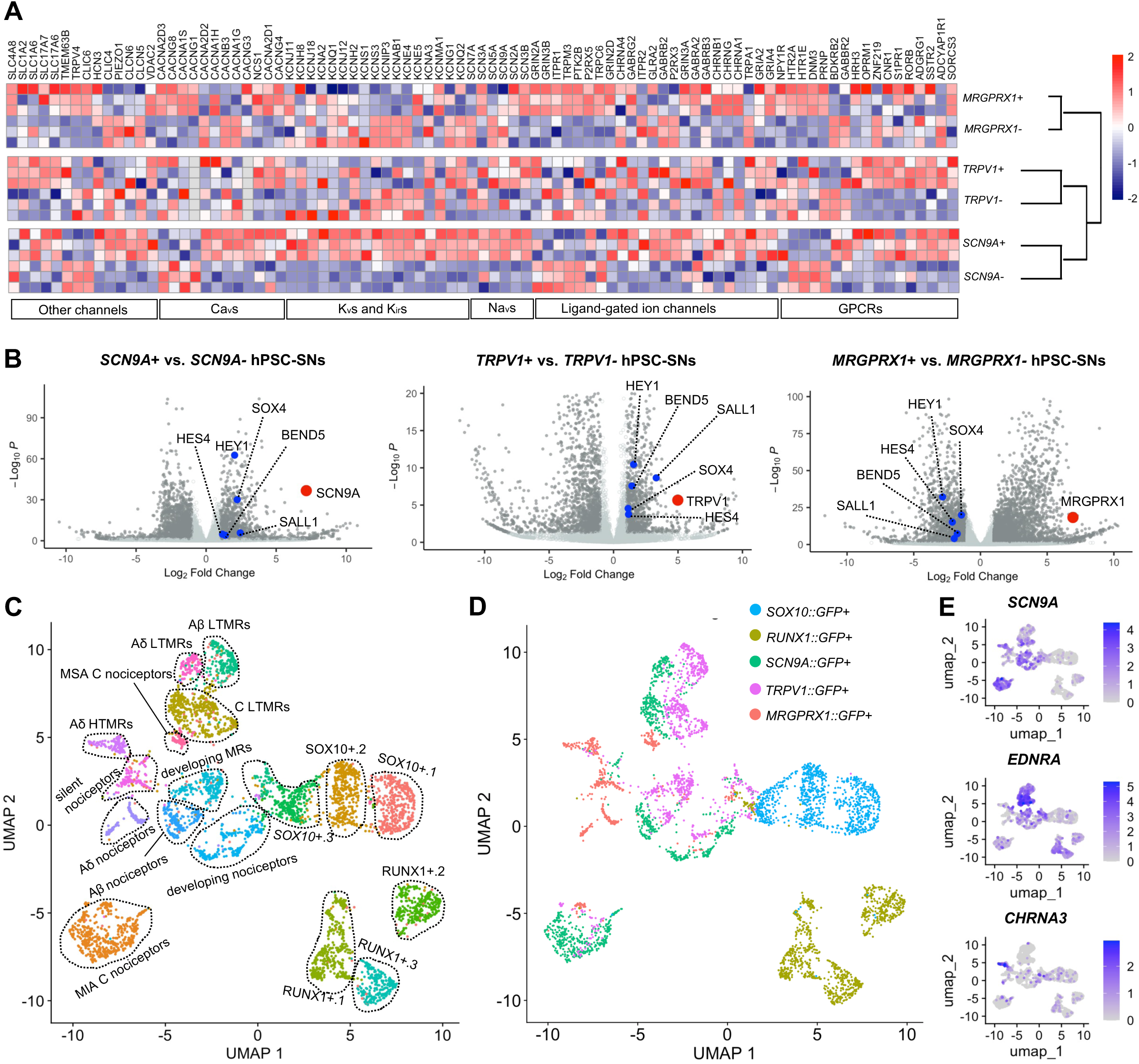
*TRPV1*+ and *SCN9A*+ nociceptors and *MRGPRX1*+ pruriceptors share transcriptomic profiles with human DRG neuron subtypes. **(A)** Gene Set Enrichment Analysis (GSEA) on differentially expressed receptors and ion channels between GFP+ and GFP-populations in hPSC-SNs from *MRGPRX1::GFP, SCN9A::GFP*, and *TRPV1::GFP* reporter lines using MSigDB gene sets; hierarchical clustering of pairwise lineage comparisons is shown as a dendrogram. **(B)** Volcano plots highlighting transcription factors differentially enriched in each GFP+ population. *SOX4*, *HEY1*, *HES4*, *BEND5*, and *SALL1* are enriched in *SCN9A*+ and *TRPV1*+ nociceptors but downregulated in *MRGPRX1*+ pruriceptors. Red dots, enrichment of *SCN9A*, *TRPV1*, and *MRGPRX1* in their respective comparison; blue dots, enrichment of transcription factors. **(C)** UMAP embedding of combined single-cell data colored by cluster identities, with 7 nociceptor clusters (“MSA C nociceptors”, “MIA C nociceptors”, “Aδ HTMRs”, “Aδ nociceptors”, “Aβ nociceptors”, “silent nociceptors”, “developing nociceptors”), 4 mechanoreceptor subtypes (“Aδ LTMRs”, “Aβ LTMRs”, “C LTMRs”, “developing MRs”) and 6 sensory neuron progenitor clusters (“*SOX10*+.1”, “*SOX10*+.2”, “*SOX10*+.3”, “*RUNX1*+.1”, “*RUNX1*+.2”, “*RUNX1*+.3”). **(D)** UMAP embedding of combined single-cell data colored by original barcoded lineage identity. **(E)** Feature plots on the UMAP embedding showing expression of marker genes used to assign each cluster identity.

Human nociceptors and pruriceptors exhibit remarkable heterogeneity in their ion channel and receptor expression. To functionally characterize nociceptive *TRPV1*+, *SCN9A*+, and pruriceptive *MRGPRX1*+ hPSC-SNs, we isolated GFP+ neurons from hPSC-SN cultures (Figure S1A) and observed that each purified population exhibited a morphologically homogeneous profile, characterized by round somata and extensive neuronal processes (Figure 1G). Cell body diameter measurements showed that all three subtypes fell within the expected range for small- and medium-fiber nociceptors and pruriceptors (15–25 µm), with *MRGPRX1*+ neurons exhibiting the lowest size variability and smallest mean diameter, consistent with their classification as a homogeneous, non-peptidergic C-fiber population^8^ (Figure 1H). We next assessed neuronal responsiveness and subtype fidelity by monitoring calcium transients in response to six canonical pruritogenic or noxious ligands: BAM8-22, capsaicin, mustard oil, menthol, histamine, and ATP (Figures S1B–S1O). As BAM8-22 and capsaicin are selective ligands for MRGPRX1 and TRPV1, respectively^9,10^, we quantified the proportion of responsive neurons within the *MRGPRX1*+ and *TRPV1*+ populations relative to their corresponding negative counterparts. *MRGPRX1*+ hPSC-SNs displayed significantly enhanced responses to BAM8-22 when compared with *MRGPRX1*– controls (Figures 1I-1K) while *TRPV1*+ hPSC-SNs exhibited markedly increased responses to capsaicin relative to *TRPV1*– cells (Figures 1L-1N). The quantification of ligand-induced activation across all three subtypes revealed distinct profiles of responsiveness to the six stimuli: *MRGPRX1*+ and *TRPV1*+ hPSC-SNs responded most robustly to their cognate agonists, whereas *SCN9A*+ neurons were preferentially activated by histamine and capsaicin (Figure 1O). Notably, *TRPV1*+ hPSC-SNs exhibited minimal responses to menthol, a TRPM8 agonist, consistent with previous findings that TRPV1 and TRPM8 are rarely co-expressed in human sensory neurons^11^. In summary, we generated and validated purifiable nociceptive *TRPV1::GFP*, *SCN9A::GFP*, and pruriceptive *MRGPRX1::GFP* reporter hPSC-SNs, demonstrating that *TRPV1*+, *SCN9A*+, and *MRGPRX1*+ hPSC-SNs represent functionally distinct nociceptor and pruriceptor subtypes with selective responses to sensory stimuli, recapitulating the polymodality of human primary afferent neurons.

### *TRPV1*+ and *SCN9A*+ Nociceptors and *MRGPRX1*+ Pruriceptors Exhibit Transcriptomic Similarity to Human DRG Neuron Subtypes

Bulk RNA transcriptomics of subtypes of human primary sensory neurons has revealed distinct molecular profiles that would otherwise be obscured in single cell RNA sequencing (scRNA-seq) due to transcriptomic noise. To delineate the transcriptomic signatures of *TRPV1*+, *SCN9A*+, and *MRGPRX1*+ hPSC-SNs, we purified each subtype and their corresponding negative populations for bulk RNA sequencing at 50-70 DIV (Figure S2A). Comparative analysis revealed *TRPV1*+, *SCN9A*+, and *MRGPRX1*+ hPSC⍰SNs exhibit partially overlapping yet distinct molecular profiles, with *MRGPRX1*+ pruriceptors displaying the most divergent transcriptome among the three subtypes (Figure S2B). We next focused on the differential expression of receptors and ion channels implicated in human nociceptive and pruriceptive signaling. Several voltage-gated potassium channels (e.g., *KCNQ2* and *KCNG1*) and ligand-gated ion channels (e.g., *TRPA1* and *P2RX3*) were preferentially enriched in *TRPV1*+ and *SCN9A*+ nociceptors but not in *MRGPRX1*+ pruriceptors, suggesting subtype-specific roles in human nociceptor excitability and signal transduction. Conversely, *MRGPRX1*+ pruriceptors were selectively enriched for in a distinct set of G-protein–coupled receptors (GPCRs), including *PRNP, DNM3, HTR1E, HTR2A,* and *NPY1R*, as well as voltage-gated calcium channels such as *CACNG1, CACNA1S, CACNG8,* and *CACNA2D3*, indicating their specialized roles in pruriceptive signaling (Figures 2A and S2C-S2E). Beyond surface receptor differences, *TRPV1*+ and *SCN9A*+ nociceptors showed enrichment in pathways and transcription factors distinct from those observed in *MRGPRX1*+ pruriceptors (Figures 2B and S2F). These findings suggest that *MRGPRX1*+ pruriceptors arise from a transcriptionally and developmentally distinct lineage compared with *TRPV1*+ and *SCN9A*+ nociceptors.

One limitation of current hPSC-based *in vitro* models is that the resulting sensory neurons lack the transcriptomic maturity observed in human dorsal root ganglion (DRG) neurons^12^. To assess the transcriptomic maturation of *TRPV1*+, *SCN9A*+, and *MRGPRX1*+ hPSC-SNs and to evaluate their similarity to adult human DRG neurons, we constructed a transcriptional trajectory encompassing SOX10+ neural crest progenitors, RUNX1+ sensory neuron progenitors, and the three hPSC-SN subtypes. Each population was isolated at its peak marker expression and subjected to scRNA-seq (Figure S3A). Clustering of the merged dataset revealed 17 distinct clusters corresponding to stages of sensory neuron development, differentiation into subtypes, and maturation^13^, including seven nociceptor clusters, four low-threshold mechanoreceptor (LTMR) clusters and six sensory neuron progenitor clusters (Figure 2C). The cluster identities were assigned based on canonical lineage markers and genes previously reported in human DRG scRNA-seq studies^7,8,14^ (Figures 2E and S3B). Notably, nearly all *TRPV1*+, *SCN9A*+, and *MRGPRX1*+ hPSC-SNs were localized within the 11 sensory neuron-committed clusters (Figure 2D). These 11 clusters, which include developing mechanoreceptors, developing nociceptors, mechanically insensitive afferent (MIA) C nociceptors, mechanically sensitive afferent (MSA) C nociceptors, Aδ high-threshold mechanoreceptors (HTMRs), Aδ nociceptors, Aβ nociceptors, silent nociceptors, Aδ LTMRs, Aβ LTMRs, and C LTMRs, exhibited transcriptomic profiles consistent with human DRG neuron subtypes; cluster markers were cross-validated against the human DRG single-cell atlas^7,8,14^ (Figures S4A-S4K). Together, bulk and single-cell transcriptomic analyses revealed distinct developmental programs and molecular signatures that differentiate nociceptive *TRPV1*+ and *SCN9A*+ hPSC-SNs from pruriceptive *MRGPRX1*+ hPSC-SNs, each closely recapitulating human DRG neuron subtypes.

### *CD200^high^* Marks a Population of hPSC-NNs

We demonstrated that nociceptive *TRPV1*+ and *SCN9A*+ hPSC-SNs closely resemble human nociceptors at both functional and transcriptomic levels because they respond to canonical noxious stimuli and express a comprehensive repertoire of nociception-related receptors and ion channels. These features led us to explore their potential as therapeutic decoys that are capable of competitively binding and sequestering local inflammatory mediators. Traditionally, tropomyosin receptor kinase A (TRKA) has been the gold standard surface marker for isolating nociceptive neurons from heterogeneous peripheral neuron populations; however, this approach is limited by its expression in a subset of non-nociceptive neurons and its absence in many bona fide nociceptors^8^. To develop a non-genetic purification strategy, we identified *CD200* as a surface marker significantly enriched in *TRPV1*+ and *SCN9A*+ nociceptors but not in *MRGPRX1*+ pruriceptors (Figures 3A and S3C). Analysis of published human DRG datasets further confirmed that *CD200* was predominantly expressed in small-diameter nociceptive afferents but not in *NEFH*+ large-diameter, non-nociceptive neurons^8^ (Figure S3D). Furthermore, single-cell transcriptomic analyses revealed the enrichment in *CD200*+ cells of pathways related to G protein signaling and anti-inflammatory cytokine production (Figure 3B) and nociceptor-specific genes and receptors for inflammatory ligands, including *SCN3B, GRIA1, GRIN2B,* and *CXCR4* (Figure 3C), both of which are critical in nociceptor inflammatory responses^15,16^. Flow cytometry confirmed that *CD200* was significantly enriched in *SCN9A*+ or *TRPV1*+ nociceptors compared with unpurified hPSC-SNs (Figures 3D and 3E). Functional characterization using calcium imaging showed that *CD200^high^* hPSC-NNs displayed heightened responses to noxious stimuli (ATP, mustard oil, capsaicin, and histamine) compared with *CD200^low^*hPSC-SNs, and responses to the pruritogen BAM8-22 and menthol were comparable (Figures 3F and S3E). Together, these results established *CD200* as a robust, non-genetic surface marker for isolating functionally validated hPSC-NNs to enable their purification for downstream therapeutic applications.

**Figure 3.**
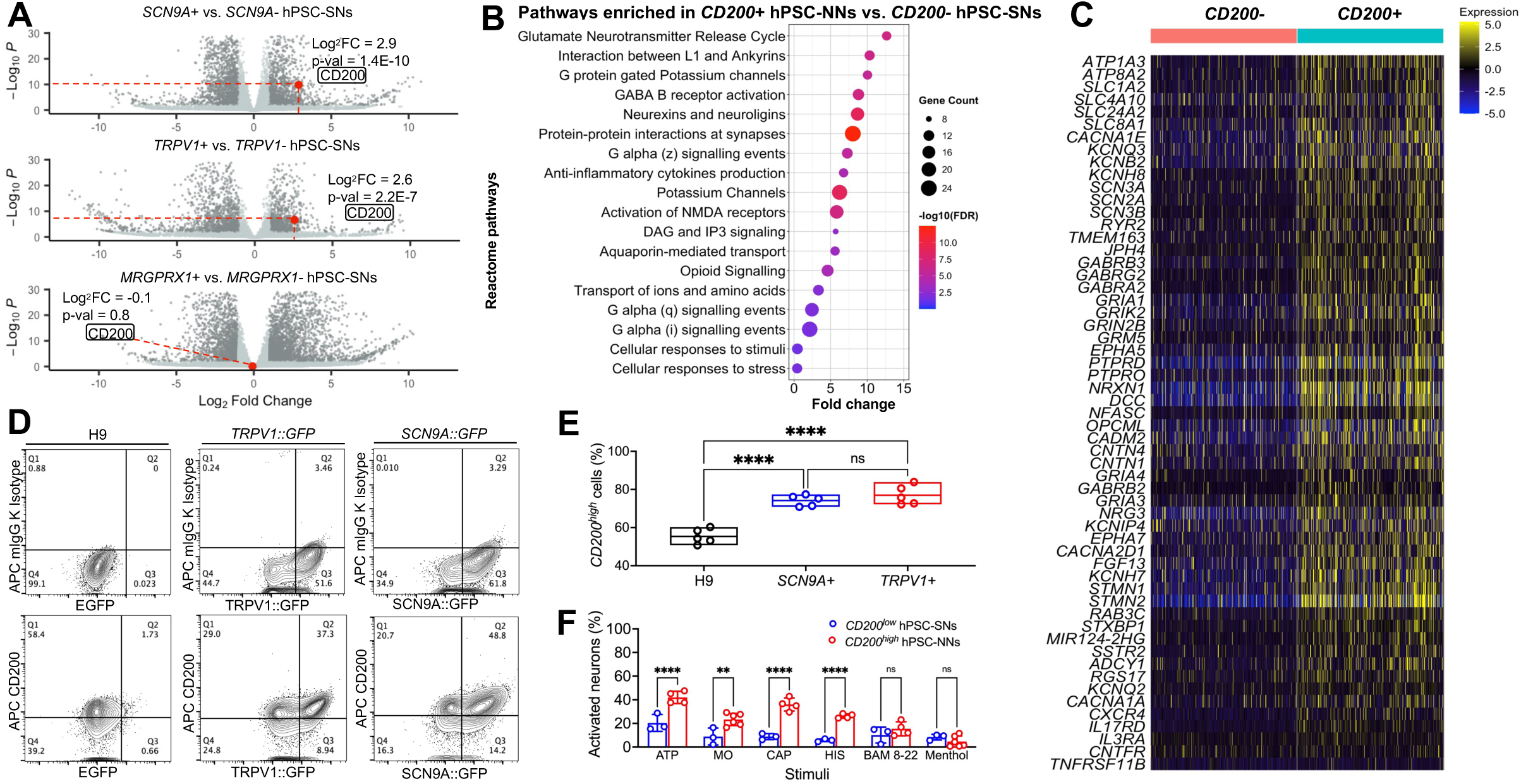
*CD200* is specifically enriched in nociceptors. **(A)** Volcano plot of bulk RNA-seq DEGs in GFP+ versus GFP-cells from *SCN9A::GFP*, *TRPV1::GFP* and *MRGPRX1::GFP* lines, highlighting enrichment of *CD200* in *SCN9A*+ and *TRPV1*+ nociceptors but not *MRGPRX1*+ pruriceptors. **(B)** Reactome pathway enrichment analysis of DEGs upregulated in *CD200*+ hPSC-NNs, showing significant enrichment of nociceptive signaling pathways. **(C)** Heatmap of z-scored expression for nociceptor-specific genes and receptors for inflammatory ligands in *CD200*+ hPSC-NNs versus *CD200-* hPSC-SNs. **(D)** FACS analysis showing the expression of *CD200* in hPSC-SNs from H9 parent line*, SCN9A::GFP*, and *TRPV1::GFP* lines. **(E)** Quantification of *CD200^high^* hPSC-NN proportions (n = 5 independent differentiations; one-way ANOVA with Tukey’s multiple comparisons test; **** P < 0.0001; mean ± range with individual values overlaid). **(F)** Bar chart showing enhanced responses of *CD200^high^* hPSC-NNs versus *CD200^low^* hPSC-SNs to ATP (50μM), mustard oil (100μM), capsaicin (10μM), and histamine (50μM) (n ≥ 3 independent differentiations; Šídák’s multiple comparisons test; ** P < 0.01, **** P < 0.0001; mean ± s.e.m. with individual values overlaid). Responses to BAM8-22 (10μM) and menthol (100μM) did not differ.

### *CD200^high^* hPSC-NNs Mitigate Pain and Modulate the Neuro-Immune Interface to Reduce Inflammation in OA Mice

Given that *CD200^high^* hPSC-NNs expressed key receptors and ion channels involved in nociceptive transduction and exhibited robust *in vitro* responses to canonical noxious stimuli, we next investigated their therapeutic potential as decoys to bind and deplete inflammatory mediators in a mouse model of OA. Forty thousand *CD200^high^*hPSC-NNs were injected into the intra-articular space and tibial tuberosity of immunocompromised NIH-III mice two weeks after bilateral anterior cruciate ligament transection (ACLT) surgery (ACLT-*CD200^high^* group)^17^. Control animals received *CD200^low^*hPSC-SNs following ACLT (ACLT-*CD200^low^* group). A sham-operated group received *CD200^low^* hPSC-SNs (sham-*CD200^low^*group) (Figure 4A). Survival of the transplanted cells within the knee joint was confirmed by immunostaining for the human-specific cytoplasmic marker STEM121 at 10 weeks post-injection^18^ (Figure 4B) with *CD200^high^* hPSC-NNs exhibiting small diameters compared with *CD200^low^* hPSC-SNs (Figure 4C). Functionally, ACLT-*CD200^high^* animals displayed significantly reduced mechanical hypersensitivity, as indicated by lower paw withdrawal frequencies in von Frey testing and elevated pressure application measurement withdrawal threshold (PAMWT) at five and 10 weeks post-injection compared with ACLT-*CD200^low^* animals (Figures 4D and 4E). Activating transcription factor 3 (ATF3) is a well-established marker of neuronal activation or stress following peripheral noxious stimulation or injury, whereas calcitonin gene-related peptide (CGRP) levels are significantly upregulated in chronic inflammatory joint pain^19–21^. To further validate the analgesic effects of *CD200^high^* hPSC-NNs, bilateral L3-L4 DRGs, which innervate the knee joint, were harvested five and 10 weeks post-injection for immunohistochemical analysis. DRGs from ACLT-*CD200^high^*animals expressed significantly decreased levels of ATF3 and CGRP compared with ACLT-*CD200^low^* animals, consistent with diminished nociceptive activity at the cellular level (Figures 4F-4H). In early OA, inflammatory signals trigger osteoclast hyperactivity, leading to subchondral bone loss whereas mid- to late-stage disease is characterized by aberrant osteoblastic differentiation, disrupting osteoclast–osteoblast coupling and increasing subchondral bone mass^2^. To assess the impact of *CD200^high^* hPSC-NNs on subchondral bone architecture, we performed 3D micro–computed tomography (µCT) analysis^22^ (Figure 4I). Bone volume fraction (BV/TV), which quantifies the proportion of mineralized bone tissue within the total volume^23^, was significantly higher at early-stage OA (five weeks) and significantly lower at late-stage OA (10 weeks) in ACLT-*CD200^high^* animals compared with ACLT-*CD200^low^*controls, maintaining subchondral bone volume comparable to sham-operated mice at both time points (Figure 4K). To assess the integrity of articular cartilage, we evaluated the Osteoarthritis Research Society International (OARSI) histopathology score 10 weeks post injection, a well validated and widely used metric for quantifying cartilage degeneration in preclinical OA models. Analysis using this scoring system revealed that *CD200^high^* hPSC-NNs significantly preserved articular cartilage and prevented degeneration compared to *CD200^low^* controls (Figures 4J and 4L). Together, these findings suggest that *CD200^high^* hPSC-NNs mitigate OA-associated pain while preserving subchondral bone architecture and protecting against cartilage degeneration in OA mice.

**Figure 4.**
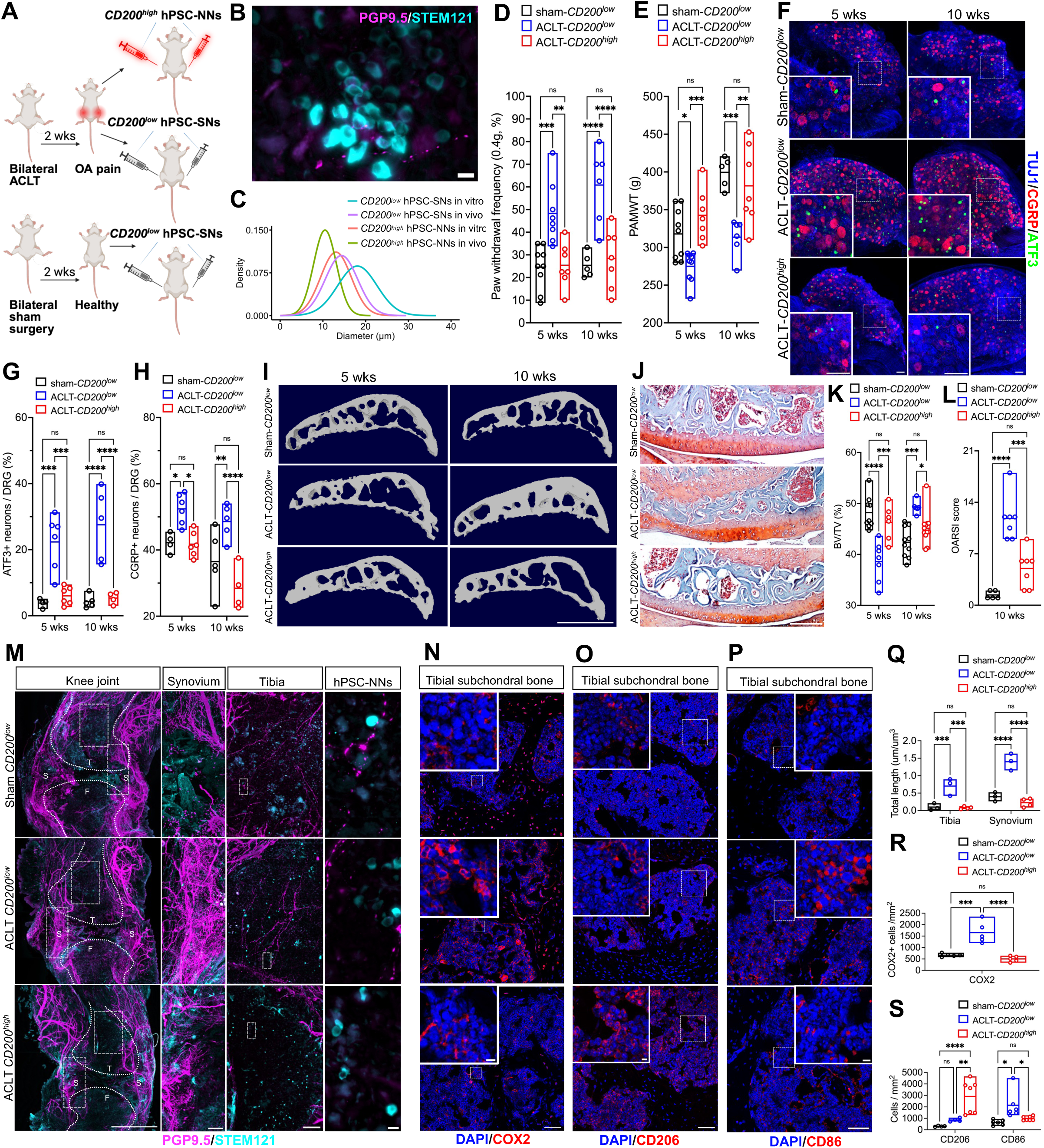
*CD200^high^* hPSC-NNs mitigate pain and modulate the neuro-immune interface to suppress inflammation in OA mice. **(A)** Experimental timeline of bilateral intra-articular and tibia tuberosity injections of *CD200^high^* hPSC-NNs or *CD200^low^* hPSC-SNs in NIH-III mice two weeks after sham or ACLT surgery. **(B)** Representative image of survival of transplanted cells in the knee joint at 10 weeks, visualized by STEM121 (cyan) and PGP9.5 (magenta) immunostaining. Scale bar, 10 μm. **(C)** Gaussian distributions of soma diameters for *CD200^high^*hPSC-NNs and *CD200^low^* hPSC-SNs measured in vitro and in vivo (n = 100 cells per group). **(D-E)** Mechanical hypersensitivity assessed by Von Frey **(D)** and PAMWT **(E)** in sham-*CD200^low^*, ACLT-*CD200^low^* and ACLT-*CD200^high^* animals 5 and 10 weeks post-injection (n ≥ 5; two-way ANOVA with Tukey’s multiple comparisons test; * P < 0.05, ** P < 0.01, *** P < 0.001, **** P < 0.0001; mean ± range with individual values overlaid). **(F-H)** L3–L4 whole DRGs immunostained for ATF3 (green) and CGRP (red) at 5 and 10 weeks; scale bars, 100µm. Quantification of ATF3+ **(G)** and CGRP+ **(H)** neurons in whole DRGs (n ≥ 4; two-way ANOVA with Tukey’s test; * P < 0.05, ** P < 0.01, *** P < 0.001, **** P < 0.0001; mean ± range with individual values overlaid). **(I)** Representative microCT of tibia subchondral bone sagittal view. Scale bar, 1mm. **(J)** Safranin O–Fast Green staining of tibial cartilage in sagittal sections. Scale bar, 200µm. **(K)** Quantification of trabecular bone volume fraction (BV/TV) (n ≥ 5; two-way ANOVA with Tukey’s test; * P < 0.05, *** P < 0.001, **** P < 0.0001; mean ± range with individual values overlaid). **(L)** Osteoarthritis Research Society International (OARSI) scores quantifying cartilage degeneration in OA (n ≥ 5; one-way ANOVA with Tukey’s multiple comparisons test; * P < 0.05, *** P < 0.001, **** P < 0.0001; data are mean ± range with individual values overlaid). **(M,Q)** 3D light-sheet microscopy of knee joints showing PGP9.5+ nerves (magenta) and STEM121+ transplanted cells (cyan) in synovium and tibia; scale bars: 1mm (far left), 100µm (left), 100µm (right), 10µm (far right). T, tibia; F, femur; S, synovium. Nerve density quantification in tibia and synovium **(o)** (n ≥ 3; two-way ANOVA with Tukey’s test; *** P < 0.001, **** P < 0.0001; mean ± range with individual values overlaid). **(N-P)** Tibial subchondral bone immunostained for COX2, CD206 and CD86; scale bars, 100 μm and 10 μm (inset). **(R-S)** Quantification of COX2, CD206 and CD86 expression (n ≥ 4; one-way ANOVA with Tukey’s test; * P < 0.05, ** P < 0.01, *** P < 0.001, **** P < 0.0001; mean ± range with individual values overlaid).

Both the nervous and immune systems play pivotal roles in the progression of OA. In particular, activated macrophages and the release of pro-inflammatory mediators promote aberrant sensory nerve sprouting, thereby amplifying articular cartilage degeneration, inflammation, and chronic pain sensitization^23–25^. To evaluate whether *CD200^high^*hPSC-NNs can modulate this pathological neuro-immune axis, we first assessed joint innervation using sectioned histology and three-dimensional whole-joint nerve mapping. Remarkably, the quantification of total nerve fiber length revealed significantly reduced innervation in the synovium and tibial subchondral bone of ACLT-*CD200^high^* animals compared with ACLT-*CD200^low^* animals at five weeks (Figures S5A and S5B) and 10 weeks post-injection (Figures 4M, 4Q, S5A and S5B), correlating with the improved behavioral pain outcomes. Given that aberrant nerve growth can be driven by elevated levels of pro-inflammatory cytokines such as COX2 and TNF-α^25^, we next examined macrophage polarization and inflammatory mediator abundance in the tibial subchondral region at 10 weeks. Immunohistochemical analysis revealed a significantly higher proportion of M2-polarized, anti-inflammatory macrophages and a marked reduction in COX2 and TNF-α expression in ACLT-*CD200^high^* animals relative to ACLT-*CD200^low^* controls (Figures 4N-4P, 4R, 4S, S5C, and S5E). Additionally, expression of matrix metalloproteinase 13 (MMP13), a mediator of cartilage degradation, was also reduced, consistent with an anti-inflammatory tissue environment (Figures S5D and S5F). To control for non-specific immunogenic or injection-related effects, we repeated the experiments and injected viable *CD200^high^* hPSC-NNs into ACLT animals (ACLT-*CD200^high^*) while using heat-killed hPSC-SNs as controls in ACLT (ACLT-heat killed) and sham-operated (sham-heat killed) groups (Figure S6A). Consistent with prior findings, *CD200^high^* hPSC-NNs conferred analgesic effects (Figures S6B-S6D, S6F, and S6G), suppressed aberrant nerve growth (Figures S6E and S6H), and preserved subchondral bone microarchitecture (Figures S6I-S6K). Together, these findings demonstrated that *CD200^high^* hPSC-NNs alleviated OA-associated pain and attenuated the pathological neuro-immune loops that drive OA progression by limiting aberrant nerve growth and suppressing pro-inflammatory signaling within the joint.

### *CD200^high^* hPSC-NNs Promote Joint homeostasis by Depleting Inflammatory Factors and Secreting Reparative Factors in Human and Mouse Joint Tissues

We next sought to determine whether the analgesic and anti-inflammatory effects of *CD200^high^* hPSC-NNs were mediated through the direct binding and sequestration of inflammatory mediators. To assess whether *CD200^high^* hPSC-NNs were activated *in vivo* in response to inflammatory stimuli, we performed immunostaining for the neuron-specific injury marker ATF3 in STEM121+ transplanted cells. Notably, *CD200^high^*hPSC-NNs displayed significantly elevated ATF3 expression compared with *CD200^low^*hPSC-SNs in ACLT animals, consistent with the hypothesis that *CD200^high^*hPSC-NNs are activated *in vivo* by endogenous inflammatory mediators (Figures 5A and 5C). To further test their responsiveness to inflammatory cues, we performed calcium imaging following acute exposure to synovial fluid from OA patients. *CD200^high^*hPSC-NNs exhibited rapid and robust calcium transients upon stimulation with synovial fluid from two independent OA donors, whereas *CD200^low^*hPSC-SNs showed delayed and significantly attenuated responses with a significantly lower proportion of responsive cells, comparable to responses elicited by pooled synovial fluid from healthy donors (Figures 5B and 5D).

**Figure 5.**
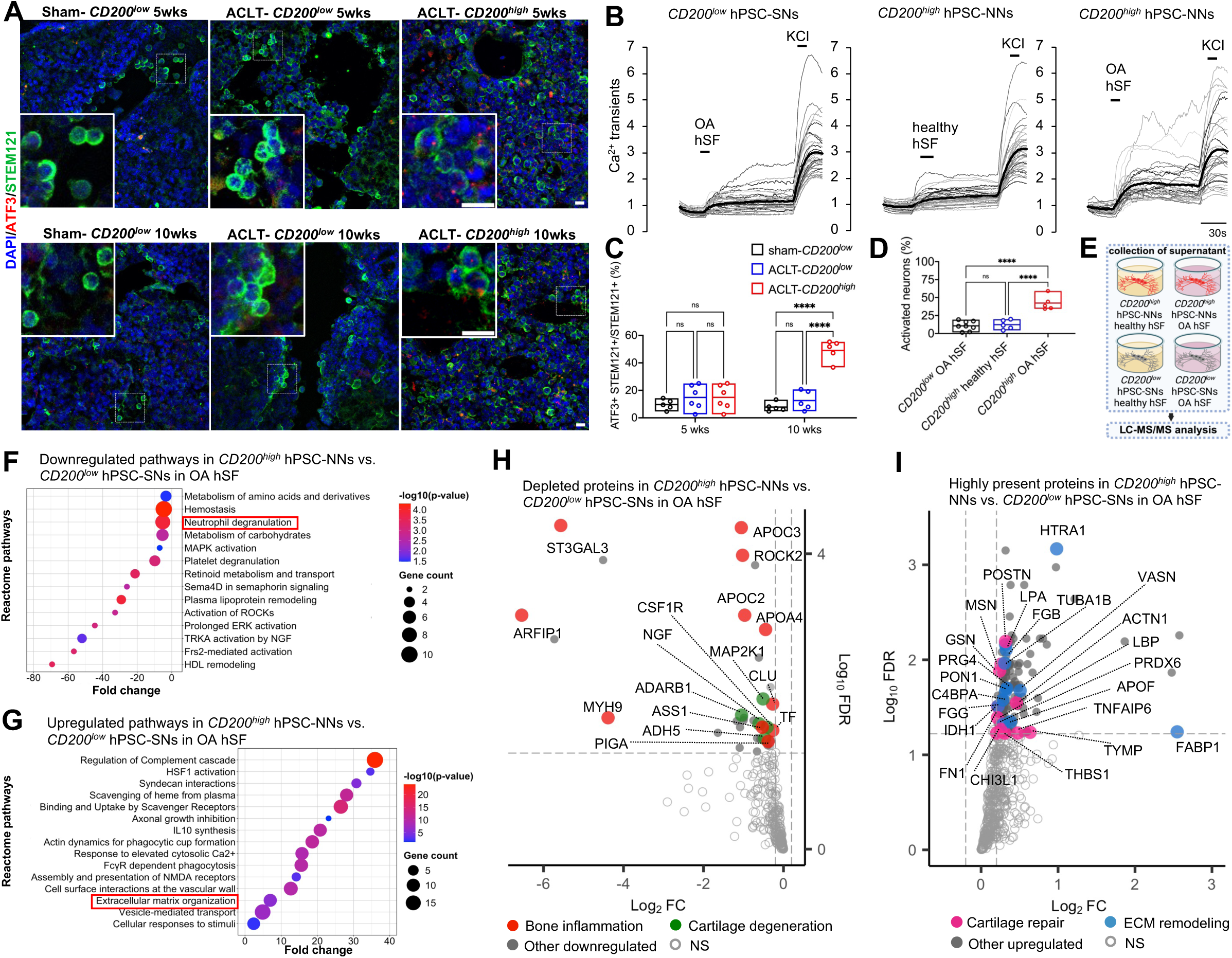
*CD200^high^* hPSC-NNs promote healthy joint homeostasis by depleting inflammatory factors and secreting factors that facilitate tissue repair. **(A,C)** Expression of ATF3 (red) in transplanted *CD200^low^* hPSC-SNs and *CD200^high^*hPSC-NNs (green) in knee sections at 5 and 10 weeks post-injection; scale bar, 10µm. Quantification of the proportion of *CD200^low^* hPSC-SNs or *CD200^high^* hPSC-NNs expressing ATF3 **(c)** (n ≥ 5; two-way ANOVA with Tukey’s multiple comparisons test; **** P < 0.0001; mean ± range with individual values overlaid). **(B,D)** Representative Ca^2+^ transients of *CD200^low^* hPSC-SNs and *CD200^high^*hPSC-NNs acutely exposed to OA hSF or pooled healthy hSF; bold traces show the mean. Quantification of portion of responsive neurons **(d**) (n = 2 independent OA donors; n ≥ 5 independent differentiations; one-way ANOVA with Tukey’s multiple comparisons test; **** P < 0.0001; mean ± range with individual values overlaid). **(E)** Schematic workflow of proteomic analysis of supernatants from *CD200^high^* hPSC-NNs and *CD200^low^* hPSC-SNs following overnight incubation with OA hSF or pooled healthy hSF (n = 2 independent OA donors different from **b,d**; n = 3 independent differentiations; n = 6 proteomic replicates). **(F)** Reactome pathway analysis showing downregulated pathways in *CD200^high^* hPSC-NNs versus *CD200^low^* hPSC-SNs incubated in OA hSF (n = 2 independent OA donors different from **b,d**; n = 3 independent differentiations; n = 6 proteomic replicates; |log_2_ fold change| ≥ 0.2; FDR cutoff < 0.05). **(G)** Reactome pathway analysis showing upregulated pathways in *CD200^high^* hPSC-NNs versus *CD200^low^* hPSC-SNs incubated in OA hSF (n = 2 independent OA donors different from **b,d**; n = 3 independent differentiations; n = 6 proteomic replicates; |log_2_ fold change| ≥ 0.2; FDR cutoff < 0.05). **(H)** Volcano plots of proteins depleted in OA hSF following incubation with *CD200^high^* hPSC-NNs versus *CD200^low^* hPSC-SNs (n = 2 independent OA donors different from **b,d**; n = 3 independent differentiations; n = 6 proteomic replicates; |log_2_ fold change| ≥ 0.2; FDR cutoff < 0.05). **(I)** Volcano plots of proteins enriched in OA hSF following incubation with *CD200^high^* hPSC-NNs versus *CD200^low^* hPSC-SNs (n = 2 independent OA donors different from **b,d**; n = 3 independent differentiations; n = 6 proteomic replicates; |log_2_ fold change| ≥ 0.2; FDR cutoff < 0.05).

To further elucidate the mechanisms underlying the analgesic and immunomodulatory effects of *CD200^high^* hPSC-NNs, we performed proteomic analysis of conditioned supernatants collected from *CD200^high^*hPSC-NNs and *CD200^low^* hPSC-SNs following overnight incubation with synovial fluid from two osteoarthritic patients (OA hSF) or pooled healthy donors (healthy hSF) (Figure 5E). Differential expression and gene ontology analysis of the proteomic profile revealed a significant downregulation of proteins implicated in bone inflammation and cartilage degeneration in OA hSF incubated with *CD200^high^* hPSC-NNs compared with *CD200^low^* hPSC-SNs. Among inflammation-associated proteins, ST3 beta-galactoside alpha-2,3-sialyltransferase 3 (ST3GAL3) activates the Toll-like receptor 9 (TLR9)/MyD88 signaling pathway^30^, whereas the apolipoproteins apolipoprotein C3 (APOC3) and apolipoprotein C2 (APOC2) promote inflammasome activation and pro-inflammatory macrophage polarization, respectively^26,27^. Among the proteins implicated in cartilage degeneration, Rho-associated coiled-coil kinase 2 (ROCK2) and mitogen-activated protein kinase kinase 1 (MAP2K1), two essential proteins in the Rho/ROCK and MEK/ERK pathways, mediate degradation of chondrocytes and the surrounding extracellular matrix (ECM)^28^. Nerve growth factor (NGF) stimulates the production of MMPs, facilitating neurovascular invasion^29,30^ (Figures 5F and 5H). Conversely, proteins and pathways implicated in cartilage repair and ECM remodeling were markedly enriched in the OA hSF group incubated with *CD200^high^*hPSC-NNs compared with *CD200^low^* hPSC-SNs. Notably, thrombospondin-1 (THBS1) exerted pro-chondrogenic effects^31^ while chitinase 3 like 1 (CHI3L1) inhibited cartilage matrix degeneration by modulating the TLR4–MAPK–STAT1 pathway^32^ (Figures 5G and 5I).

To control for passive release of intracellular contents from dead cells, we conducted proteomic analysis on knee joints harvested 10 weeks post-injection from sham-heat killed, ACLT-heat killed, and ACLT-*CD200^high^*groups to identify differentially regulated mouse proteins (Figure 6A). To eliminate potential cross-species contamination, precursors mapping to both human and mouse proteins were excluded, and only mouse-specific precursors were retained for quantification. Quantification of differentially expressed proteins across the three groups revealed a divergence of the proteomes between the ACLT-heat killed and sham-heat killed groups, and a remarkable similarity between the proteomes of ACLT-*CD200^high^* and sham-heat killed animals (Figure 6B). Differential expression and gene ontology analysis showed factors implicated in inflammation and joint degeneration were significantly enriched in ACLT-heat killed animals but, strikingly, were reversed in ACLT-*CD200^high^* animals. Among these proteins, matrix metalloproteinase-8 (Mmp8) promotes chondrocyte apoptosis and cartilage degradation^33^, and mitogen-activated protein kinase 13 (Mapk13) increases expression of catabolic enzymes^34^ (Figures 6D, 6F, and 6G). Conversely, proteins and pathways involved in anti-inflammation, ECM remodeling, and joint homeostasis were suppressed in ACLT-heat killed animals but restored in ACLT-*CD200^high^*animals. Among these proteins, olfactomedin 1 (Olfm1) inhibits osteoclast differentiation^35^, while ECM components including matrilin-1 (Matn1), matrilin-1 (Matn3), tenascin XB (Tnxb), and multiple collagen peptides are essential for cartilage integrity^36^(Figures 6E, 6H, and 6I). Notably, among the genes enriched in both OA hSF and mouse knee joints following *CD200^high^* hPSC-NN treatment, paraoxonase 1 (Pon1) is an enzyme that protects lipoproteins from oxidative modifications^37^. These findings suggest that *CD200^high^* hPSC-NNs secrete factors that stimulate host tissues to produce pro-reparative proteins. Together, these data suggest that *CD200^high^* hPSC-NNs act through a dual mechanism in OA, simultaneously sequestering pro-inflammatory mediators and secreting factors that promote bone and ECM remodeling, thereby creating a reparative, anti-inflammatory microenvironment in human and mouse joint tissues (Figure 6C).

**Figure 6.**
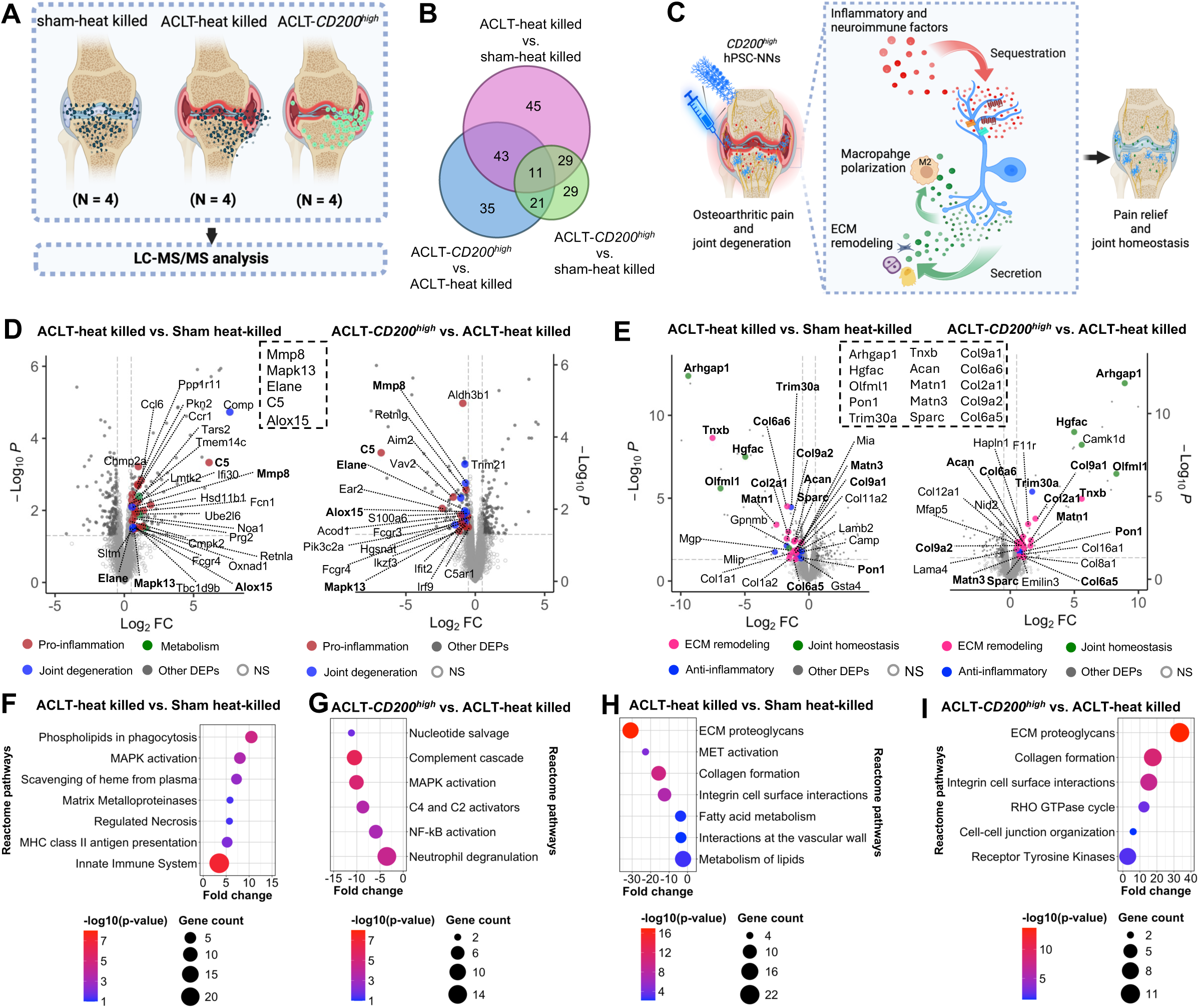
*CD200^high^*hPSC-NNs restored the disrupted proteome in OA mice, promoting an anti-inflammatory and reparative microenvironment. **(A)** Schematic workflow of proteomic analysis of knee joints harvested 10 weeks post-injection from ACLT animals injected with viable *CD200^high^* hPSC-NNs (ACLT-*CD200^high^*), ACLT animals injected with heat-killed hPSC-SNs (ACLT-heat killed), and sham-operated animals injected with heat-killed hPSC-SNs (sham-heat killed). **(B)** Venn diagram shows differentially expressed proteins, highlighting similarity between the proteomes of sham-heat killed and ACLT-*CD200^high^* animals (n = 4 animals; n = 4 proteomic replicates; log_₂_FC cutoff = 2SD; P cutoff = 0.05). **(C)** Schematic representation of the dual mechanism of action of *CD200^high^* hPSC-NNs in OA mice. **(D)** Volcano plots of proteins enriched in ACLT-heat killed versus sham-heat killed animals (left) and reversed by *CD200^high^* hPSC-NNs 10 weeks post-injection (right); shared proteins are boxed (n = 4 animals; n = 4 proteomic replicates; |log_2_ fold change| ≥ 0.5; P cutoff = 0.05). **(E)** Volcano plots of proteins downregulated in ACLT-heat killed versus sham-heat killed animals (left) and restored by *CD200^high^* hPSC-NNs 10 weeks post-injection (right); shared proteins are boxed (n = 4 animals; n = 4 proteomic replicates; |log_2_ fold change| ≥ 0.5; P cutoff = 0.05). **(F,G)** Reactome pathways enriched in ACLT-heat killed versus sham-heat killed animals **(F)** and reversed by *CD200^high^* hPSC-NNs 10 weeks post-injection **(G)** (n = 4 animals; n = 4 proteomic replicates; |log_2_ fold change| ≥ 0.5; P cutoff = 0.05). **(H,I)** Reactome pathways downregulated in ACLT-heat killed versus sham-heat killed animals **(H)** and restored by *CD200^high^* hPSC-NNs 10 weeks post-injection **(I)** (n = 4 animals; n = 4 proteomic replicates; |log_2_ fold change| ≥ 0.5; P cutoff = 0.05).

## DISCUSSION

hPSC-derived cells have long been explored for cell replacement applications across diverse diseases such as neurological disorders, diabetes, musculoskeletal injury, and hepatic or renal dysfunction because of their renewal and pluripotency properties^38^. Here, we present an alternative application of hPSC-derived cells that diverges from this traditional regenerative paradigm. Through DECIR, we demonstrate that ectopically transplanted hPSC-NNs can function as therapeutic agents by modulating the inflammatory microenvironment through both ligand sequestration and the release of reparative factors in mouse and human joint tissues.

A major barrier to both mechanistic studies and translational advancement in pain research has been the inability to isolate nociceptor subtypes from heterogeneous sensory neuron cultures. To overcome this, we developed both genetic and non-genetic strategies to purify molecularly defined hPSC-derived nociceptors. First, we engineered GFP-reporter hPSC lines targeting *TRPV1*, *SCN9A*, and *MRGPRX1*, enabling the isolation of nociceptive (*TRPV1*+ and *SCN9A*+) and pruriceptive (*MRGPRX1*+) subtypes. Functional, morphological, and transcriptomic analyses confirmed that these purified neurons closely recapitulate their native human counterparts, providing a powerful platform for high-throughput drug screening and target discovery in pain and itch research. However, the use of these genetically labeled cells in cell therapy is limited due to concerns about the immunogenicity and cytotoxicity of fluorescent proteins^39^. To allow non-genetic, clinically actionable purification of hPSC-NNs, we identified *CD200* as a nociceptor-specific surface marker enriched in *TRPV1*+ and *SCN9A*+ nociceptors. *CD200^high^* hPSC-NNs respond robustly to inflammatory mediators in synovial fluid from OA patients and are transcriptomically enriched for ion channels, receptors, and pathways essential for nociceptive signaling.

Importantly, in the adult peripheral nervous system, the capacity for *de novo* neurogenesis under physiological conditions is minimal^40^. As such, transplantation of hPSC-NNs at quantities vastly exceeding endogenous nociceptor populations offers a competitive advantage for binding and depleting inflammatory factors. In this setting, DECIR therapy intercepts noxious signals before they engage endogenous nociceptive circuits, thereby reducing chronic pain and modulating the joint microenvironment. As a multifaceted disease, OA is characterized by a self-perpetuating cycle involving the nervous system, immune responses, and uncoupled bone remodeling processes. This intricate interplay contributes to both the progression of joint degeneration and the persistence of chronic pain. Clinically, OA often first manifests as synovial inflammation and pain, followed by microenvironmental alterations in subchondral bone and progressive structural changes in both cartilage and bone tissue. Consequently, early-stage OA management primarily targets pain control and inflammation suppression. When injected into the knee joints of OA mice, *CD200^high^* hPSC-NNs serve as scavengers of pro-inflammatory mediators. These cells alleviate OA-associated pain and modulate the endogenous neuro-immune interface by reducing aberrant nerve growth and suppressing pro-inflammatory signaling, thereby disrupting the vicious cycle that fuels OA progression. The ability of *CD200^high^*hPSC-NNs to maintain tibial subchondral bone mass in OA animals to levels comparable to the control group may stem from their capacity to disrupt interoceptive signaling between the knee joint microenvironment and the sensory nervous system^41^. This suggests a potential neuro-modulatory role for *CD200^high^* hPSC-NNs in attenuating pathological feedback loops that drive disease progression.

Despite the promising efficacy of *CD200^high^* hPSC-NNs in our OA mouse model, several important considerations remain regarding their translational potential. First, although these cells are terminally differentiated, comprehensive long-term tumorigenicity studies are needed to confirm their *in vivo* safety. Second, anatomical and volumetric differences between murine and human knee joints present challenges for clinical translation, requiring careful optimization of dosing, delivery route, and cell persistence. Addressing these translational hurdles will be critical for advancing *CD200^high^* hPSC-NNs as a viable cell-based therapy for OA and other chronic inflammatory joint diseases.

Beyond advancing hPSC-based therapies, our study uncovers a previously unrecognized reparative function of nociceptors in response to environmental insults. While the secretory profiles of neural stem cells have been well characterized^42^, our proteomic analyses reveal that hPSC-derived nociceptors secrete a distinct reparative secretome that promotes regeneration of adjacent tissues. The signaling pathways that trigger this inflammation-induced secretome remain unknown and will be the subject of future research. Notably, a recent study demonstrated that hPSC-derived sensory neurons activate phosphorylated signal transducer and activator of transcription 3 (pSTAT3) in response to rheumatoid arthritis synovial fluid^43^, suggesting that STAT3 activation, alongside other pathways, may contribute to the induction of this reparative program.

In summary, we demonstrate that ectopically grafted *CD200^high^*hPSC-NNs act as multifunctional therapeutic agents for osteoarthritis. This conceptual advance, which we term DECIR, extends the utility of hPSC-based therapies beyond traditional tissue replacement. By leveraging *CD200* as a clinically actionable surface marker, we achieve selective enrichment of hPSC-NNs capable of surviving and functioning in a non-native joint environment. Mechanistically, *CD200^high^* hPSC-NNs both sequester inflammatory mediators and secrete a reparative secretome, relieving pain and restoring joint homeostasis. These findings reveal an unexpected regenerative function of nociceptors in response to inflammatory challenge. This work redefines the therapeutic potential of hPSC-derived neurons and paves the way for their clinical translation in treating complex neuroimmune and musculoskeletal disorders.

## MATERIALS AND METHODS

### Maintenance of hPSCs

Healthy control hESC (H9, WiCell) and hiPSCs (KOLF2.1J) were cultured using standard protocols. The hPSC line was cultured with mouse embryonic fibroblasts (MEFs) (Gibco) pre-plated at 12,000–15,000 cells/cm^2^. hPSC culture medium contained DMEM/F12 (Invitrogen), 20% knockout serum replacement (Gibco), 0.1□mM MEM-NEAA (Gibco), 1□mM l-glutamine (Gibco), 55□μM β-mercaptoethanol (Gibco) and 4□ng/ml FGF2 (Gibco) with 10□μM Y-27632 (Cayman Chemical). Cells were fed daily, and passaged weekly using Accutase (Innovative Cell Technologies).

### Generation of CRISPR knock-in *TRPV1::GFP*, *SCN9A::GFP*, and *MRGPRX1::GFP* hPSC lines

Knock-in strategy using sgRNA:Cas9 ribonucleoprotein (RNP) complex via nucleofection was performed as previously described^45,46^. For each locus, a self-cleaving 2A⍰eGFP⍰PGK⍰Puro cassette was inserted into the 3′ UTR via CRISPR–Cas9-mediated homology-directed repair (HDR), with the RNP complex delivered by nucleofection. To enhance HDR efficiency and suppress non-homologous end joining (NHEJ), we co-introduced the histone deacetylase inhibitor valproic acid (VPA) and the i53 plasmid^47^. During nucleofection, H9 hESCs were dissociated into single cells using Accutase (Innovative Cell Technologies), and 1.5× 10^6^ H9 cells were resuspended in nucleofection solution V (Lonza) with 5μg HDR donor plasmid, 2 μg i53 plasmid (Addgene, #119002), 0.5 nmol sgRNA (IDT), and 0.5 nmol Cas9 nuclease (IDT, HiFi V3) to generate *TRPV1::GFP*, *MRGPRX1::GFP* and *SCN9A::GFP lines*.

Nucleofection was performed with Nucleofector™ II according to the manufacturer’s instruction (B-16 program, Lonza). The nucleofected cell suspension was subsequently plated on puromycin-resistant mouse embryonic fibroblasts (DR4, Global Stem) in hESC medium with 10 μM Y-27632 (Cayman Chemical). Three days after nucleofection, the cells that had undergone homologous recombination were selected using 0.5□μg/ml puromycin (MilliporeSigma) in hES medium for two days. After selection, puromycin-resistant colonies for each line were verified for GFP expression by FACS analysis using the differentiation protocol. n ≥ 3 clones were validated for each line. The following sgRNA and HDR template sequences were used:

*MRGPRX1* sgRNA: GGAAGCAGATTGGAGCAGTG
*MRGPRX1* HDR (left arm forward): ATAGGATCCGCCTTATATATTCCCTGTTAAGCT
*MRGPRX1* HDR (left arm reverse): ATAGCTAGCCTGCTCCAATCTGCTTCCCGACAGGC
*MRGPRX1* HDR (right arm forward): ATAGGCGCGCCGGAAGCGCCTCTGCCCTGTCAGACA
*MRGPRX1* HDR (right arm reverse): ATAGCGGCCGCCAATGGCCAAAATAGAACTGGGGCA
*TRPV1* sgRNA: GCCGCTTCCGGGGAGAAGTG
*TRPV1* HDR (left arm forward): ATAGGATCCGCTGCAAGCAAGTCGGGGTGGGA
*TRPV1* HDR (left arm reverse): ATAGCTAGCCTTCTCCCCGGAAGCGGCAGGAC
*TRPV1* HDR (right arm forward): ATAGGCGCGCCGAACGTCACGCAGACAGCACTG
*TRPV1* HDR (right arm reverse): ATAGCGGCCGCAAGCAGAAGAATCGCTTGAACCC
*SCN9A* sgRNA: AGACAAAGGGAAAGACAGCA
*SCN9A* HDR (left arm forward): ATAGGATCCGTTAGTTATATCATCATATCCTTCC
*SCN9A* HDR (left arm reverse): ATAGCTAGCTTTTTTGCTTTCCTTGCTGTCTTTC
*SCN9A* HDR (right arm forward): ATAGGCGCGCCAGCTTCATTTTTGATATATTGTTT
*SCN9A* HDR (right arm reverse): ATAGCGGCCGCCTATGAACAAAAAGGGATATGA

Nucleofection was performed with Nucleofector™ II according to the manufacturer’s instruction (B-16 program, Lonza). The nucleofected cell suspension was subsequently plated on puromycin-resistant mouse embryonic fibroblasts (DR4, Global Stem) in hESC medium with 10 μM Y-27632 (Cayman Chemical). Three days after nucleofection, the cells that had undergone homologous recombination were selected using 0.5□μg/ml puromycin (MilliporeSigma) in hES medium for two days. After selection, puromycin-resistant colonies for each line were verified for GFP expression by FACS analysis using the differentiation protocol. n ≥ 3 clones were validated for each line.

### Differentiation of hPSC-SNs

hPSCs were dissociated into single cells using TrypLE Express (Thermo Fisher Scientific) and pre-plated on 0.1% gelatin (STEMCELL Technologies) for 10 minutes to deplete mouse embryonic fibroblasts. Non-adherent hPSCs were then collected and seeded at 4.6 × 10^4^ cells/well in 24-well plates coated with 1% Geltrex™ (Gibco) in DMEM (Gibco). Cells were maintained in MEF-conditioned KSR medium (DMEM/F-12, 20% knockout serum replacement, 0.1 mM MEM-NEAA, 1 mM L-glutamine, 55 µM β-mercaptoethanol) supplemented with 10 µM Y-27632. Neural crest differentiation was induced as previously described^48^. On day 12, neural crest cells were dissociated with TrypLE Express and replated at 7 × 10^4^ cells per well on 12-well plates coated with 15 μg/ml poly-l-lysine (Sigma-Aldrich, P2636), 1 μg/ml laminin (Thermo Fisher Scientific, 217015) and 10 μg/ml fibronectin (Cell Signaling Technology, 26836SF). Sensory neuron (SN) medium was prepare using Brainphys medium (STEMCELL Technologies, # 05790) supplemented with 1% N2 supplement, 2% B27 supplement, 0.1□mM MEM-NEAA, 1□mM l-glutamine, 1% Penicillin-Streptomycin (Gibco), 20 ng/ml NGF (Sigma-Aldrich), 10 ng/ml GDNF (Sigma-Aldrich), 200 µM dibutyryl cAMP (Sigma-Aldrich) and 200 µM sodium l-ascorbate (odium l-ascorbate). Cells were suspended in SN medium as 5 µL droplets, allowed to attach for 10 min at 37 °C, then overlaid with SN medium; laminin and fibronectin were replenished weekly to maintain attachment.

### Sorting of hPSC subtypes

One day prior to FACS sorting, mature hPSC-SNs were fed with fresh SN medium. On the day of FACS purification, conditioned medium was harvested from each well, filtered, and supplemented with 10 μM of Y-27632 to make post-sorting medium (PSM). hPSC-SNs (>60-70 DIV) were incubated with TrypLE Express and DNase I for 30 minutes at 37 °C to dissociate into single cells. Digested cells were resuspended in FACS media (90% DPBS + 10% DMEM, 10 μM of Y-27632, and 3% penicillin-streptomycin) and passed through 50 um strainers (Corning). For purification of *TRPV1*+, *SCN9A*+, or *MRGPRX1*+ hPSC-SNs, 1 ul Sytox Red (ThermoFisher Scientific, S34859) was added per 1 ml cell suspension. For sorting of *CD200^high^* hPSC-NNs, 1 ul LIVE/DEAD™ Fixable Violet Dead Cell Stain (Invitrogen) and 10 ul CD200/APC (AAT Bioquest) were added per 1 ml cell suspension and incubated in the dark for 10 minutes at 37 °C. Cells were washed twice with PBS and passed through 50 um strainers again. Cells were sorted using SH800S Cell Sorter and collected in 1.5 ml Eppendorf tubes.

### Flow cytometry

Cultured hPSC-SNs (50–70 DIV) were incubated with TrypLE Express and DNase I for 30□min at 37□°C to dissociate them into single cells. The digested cells were resuspended in FACS media (90% DPBS + 10% DMEM, 10□μM Y-27632) and passed through 50□μm strainers (Corning). For flow cytometry of *TRPV1*+, *SCN9A*+, or *MRGPRX1*+ hPSC-SNs, 1□μl Sytox Red (Thermo Fisher Scientific, S34859) was added per 1□ml cell suspension. For flow cytometry of CD200, 1□μl LIVE/DEAD™ Fixable Violet Dead Cell Stain (and 10□μl CD200/APC or 10□μl Mouse IgG1 kappa Isotype Control (eBioscience) were added per 1□ml cell suspension and incubated in the dark for 10□min at 37□°C. After incubation, cells were washed twice with PBS and passed again through 50□μm strainers. Flow cytometry was performed using SH800S Cell Sorter and analyzed using FlowJo (v10.9.0).

### *In vitro* calcium imaging and analyses

Purified cells were pelleted, resuspended in PSM at a density of 1 × 10^4^ cells per 5□μl, and replated in 48-well plates pre-coated with 15□μg/ml poly-L-lysine, 1□μg/ml laminin, and 10□μg/ml fibronectin. A 5□μl droplet was placed in each well, and cells were incubated at 37□°C for 10□min to allow full attachment before adding PSM medium. Replated cells were cultured for an additional 5–7 days to ensure recovery from sorting-induced cellular stress. For calcium imaging, cells were loaded with 5□μM Calbryte™ 630 AM (AAT Bioquest) at 37□°C for 45□min in calcium imaging buffer (CIB: 10□mM Hepes, 1.2□mM NaHCO_3_, 130□mM NaCl, 3□mM KCl, 2.5□mM CaCl_2_, 0.6□mM MgCl_2_, 20□mM glucose, 20□mM sucrose; pH 7.4, 290–300□mOsm). After incubation, cells were washed in CIB to remove excess dye. Imaging was performed using a Nikon Ti2-E fluorescence microscope. For analyses of neuronal responsiveness to canonical stimuli, the following stimuli were applied: BAM8-22 (10□μM), capsaicin (10□μM), mustard oil (100□μM), menthol (100□μM), histamine (50□μM), and ATP (50□μM). KCl (100□mM) and ionomycin (1□μM) were used as positive controls for neuronal and general cell viability, respectively. For analyses of neuronal responsiveness to human synovial fluid, samples were obtained from Biotrend (normal: CDD-H-5000-N; OA: CDD-H-5000-OA). Normal synovial fluid was pooled from multiple donors, while OA synovial fluids were acquired from two independent donors. Frames were captured every 1□s for 3–4□min using a 10× objective. Calcium imaging analysis was performed with Nikon NIS-AR software. Regions of interest (ROIs) were automatically detected (single object segmentation; textural compactness = 40; object filter: 15□<□equivalent diameter□<□40), and ROIs showing no response to KCl were excluded. Neuronal responses were calculated as ΔF/F = (F_t_ – F₀)/F₀. Calcium transients were defined as chemically induced if they occurred between the start of chemical administration and up to 15□s after the end of administration. A ΔF/F ratio >□0.25 was considered a positive neuronal response.

### Immunofluorescence of hPSC-SNs

Cultured neurons were fixed with 4% PFA for 15 mins at room temperature. After 2 washes with PBS and the samples were blocked in PBS with 5% Bovine Serum Albumin (BSA) (Sigma Aldrich) and 0.2% Triton X-100 (Sigma Aldrich) for 1 hour at room temperature and incubated with primary antibodies overnight at 4°C. The following primary antibodies were used for staining: anti-TUJ1 (mouse; Abcam, ab78078; 1:800); anti-GFP (chicken; Abcam, ab13970; 1:800). Primary antibodies were washed 3 times with PBS with 0.3% Triton X-100 (PBST). The samples were incubated with secondary antibodies for 1-2 hours at room temperature. The following secondary antibodies were used for staining: Alexa Fluor 488 donkey anti-chicken IgY (H+L) (ThermoFisher Scientific; A78948; 1:800), Alexa Fluor 594 donkey anti-mouse IgG (H+L) (ThermoFisher Scientific; A-21203; 1:800). After secondary antibody incubation, cells were washed 3 times with PBST and mounted using VECTASHIELD Antifade Mounting Medium with DAPI (Fisher Vector Lab) in the culture plate and imaged using Zeiss LSM700 Confocal microscope.

### Bulk RNA-seq and analyses

Sorted cells (50-70 DIV) were collected and submitted to Johns Hopkins Transcriptomics and Deep Sequencing Core Facility for NextSeq mid output flow cell at 2×75 paired end reads. RNA⍰seq reads were first assessed for quality using FastQC and, where necessary, adapter and low⍰quality sequences were trimmed with Trim□Galore. Trimmed paired⍰end reads were aligned to the hg19 reference genome using STAR (v2.5.0), with splice⍰junction annotations supplied via GENCODE v38 GTF. Resulting coordinate-sorted BAM files were used to generate HOMER (v4.11) tag directories, from which gene-level counts were obtained by analyzeRepeats.pl. A unified count matrix was assembled across all samples, and differential expression analysis was performed in HOMER with a false-discovery rate cutoff of 0.05 and absolute log2 fold change ≥1. Downstream visualization and statistical validation, including volcano plots, pie charts, and heatmaps, were carried out in R (v4.3.2). Gene-set enrichment analysis (GSEA) was performed on *TRPV1*+ vs. *TRPV1*-, *SCN9A*+ vs. *SCN9A*- and *MRGPRX1*+ vs. MRGPRX1-populations using pairwise comparisons to assess enrichment across gene sets from MSigDB, including membrane receptor–related genes, KEGG pathways, and transcription factor targets. Analyses were run with the following parameters: FDR ≤ 5%, minimum gene set size = 10, maximum gene set size = 500, and 2,000 permutations.

### scRNA-seq and analyses

Sorted *SOX10*+, *RUNX1*+, *MRGPRX1*+, *SCN9A*+, and *TRPV1*+ cells were collected and submitted to the Johns Hopkins Transcriptomics and Deep Sequencing Core Facility for single-cell RNA sequencing (scRNA-seq) using the 10x Genomics 3′ protocol. Gel Bead-In-EMulsions (GEMs) were generated and barcoded according to the manufacturer’s instructions. To recover ∼1,000 cells per lineage, >3,000 cells were loaded onto a Chromium Next GEM Single Cell 3′ chip (10x Genomics). cDNA libraries were prepared using the Chromium Next GEM Single Cell 3′ GEM, Library & Gel Bead Kit v3.1 and sequenced on an Illumina NovaSeq S4 platform (2 × 150 bp).

Unique molecular identifier (UMI) counting was performed with Cell Ranger (v7.0.1) using GRCh38/Ensembl. Downstream analyses were conducted using R (v4.3.2) and Seurat (v5). Sequencing reads for each population were processed as separate Seurat objects and merged while retaining original barcodes. Cells with <2,500 or >10,000 detected genes or >5% mitochondrial content were excluded. Each lineage was downsampled to ∼1,000 cells. Gene expression was normalized using the LogNormalize method (scale factor = 10,000), and SCTransform was applied for variance stabilization. The top 50 principal components were used for clustering (FindNeighbors, FindClusters) and visualization with UMAP. Clusters were annotated based on canonical markers from human DRG scRNA-seq datasets.

### Mice, ACLT surgery and cell injection

All experiments were performed in accordance with protocols approved by the Animal Care and Use Committee at the Johns Hopkins University School of Medicine. We established anterior cruciate ligament transection (ACLT) model for NIH III nude mice (Charles River Laboratories) at 2 months of age. Animals were anesthetized via intraperitoneal injection of ketamine (100 mg/kg) and xylazine (15 mg/kg). A 5 mm longitudinal incision was made along the medial parapatellar region, and the patella was displaced laterally to expose the joint. Under a surgical microscope, the anterior cruciate ligament was transected with micro-scissors, the joint was irrigated with sterile saline to clear debris, and the patella was repositioned. The joint capsule, subcutaneous tissue, and skin were closed in separate layers using absorbable sutures. Sham-operated controls underwent identical exposure without ligament transection. Two weeks post-surgery, we prepared *CD200^high^*hPSC-NNs, *CD200^low^* hPSC-SNs, or heat-killed cells by resuspending them in DPBS at 1.3 × 10^6^ cells per ml. A volume of 30–40 µl of cell suspension was delivered into the intra-articular space and tibial tuberosity via a 32-gauge syringe. Mice were euthanized at 5 or 10 weeks after ACLT by transcardial perfusion with PBS (n ≥ 5 per group) for downstream analyses.

### Confocal and light sheet microscopy of mouse tissues

Mice were anesthetized and transcardially perfused with PBS followed by 10% neutral-buffered formalin. Knee joints and L3–L4 DRGs were harvested, post-fixed in 10% formalin for 24 h, and decalcified in 0.5 M EDTA (pH 7.4) at 4□°C for 14 days with gentle agitation.

#### Knee joint sections

For immunofluorescence staining and imaging of knee joints, the joints were collected and decalcified by 0.5M ethylenediaminetetraacetic acid (EDTA, pH7.4) for two weeks as previously reported^49^. Samples were then cryoprotected in 30% sucrose, embedded in OCT, and frozen. Sagittal sections (40□μm) were cut on a cryostat and mounted onto charged glass slides. Sections were permeabilized in PBS + 0.3% Triton X-100 and blocked in 5% normal donkey serum for 1□h at room temperature. Tissues were incubated at 4□°C for 48□h with the following primary antibodies in blocking buffer: anti-CGRP (goat; Abcam, ab36001, 1:200), anti-PGP9.5 (rabbit; Sigma, SAB4503057, 1:400), anti-CD206 (rabbit; Abcam, ab64693, 1:500), anti-CD86 (rat; BD Biosciences, 553689, 1:100), anti-COX2 (rabbit; Cell Signaling, 12282, 1:200), anti-MMP13 (rabbit; Abcam, ab39012, 1:400), anti-TNFα (rabbit; Proteintech, 17590-1-AP, 1:500), and anti-STEM121 (mouse; Takara, Y40410, 1:200). After three washes with PBST, sections were incubated for 1□h at room temperature in the dark with the following secondary antibodies: Alexa Fluor 488 Donkey anti-Mouse IgG (Thermo Fisher Scientific, A-21202, 1:500), Alexa Fluor 647 Donkey anti-Goat IgG (Thermo Fisher Scientific, A-21447, 1:500), Alexa Fluor 594 Donkey anti-Rabbit IgG (Thermo Fisher Scientific, A-21207, 1:500), and DAPI (1:250; Vector Laboratories, H-1200). Imaging was performed using a Zeiss LSM 880 confocal microscope, and quantification was carried out in Fiji.

#### Whole DRGs

For immunofluorescence staining and imaging of whole mouse DRGs, fixed DRGs (2–4 per condition) were incubated in at least 1□ml permeabilization solution (40□ml PBS, 0.2% Triton X-100, 1.15□g glycine, and 10□ml DMSO) for 24□h at 37□°C. Samples were then washed three times in PBS + 0.5% Triton X-100 and incubated in 1□ml blocking solution (10% normal donkey serum in PBS + 0.5% Triton X-100) for 4□h at room temperature or overnight at 4□°C. For primary antibody staining, samples were incubated in 0.5–1□ml primary antibody solution (primary antibody diluted in 5% normal donkey serum, 1% BSA in PBS + 0.5% Triton X-100) overnight at 37□°C and then for 48□h at 4□°C. The following primary antibodies were used: anti-ATF3 (rabbit; Invitrogen, PA5-101089, 1:400), anti-CGRP (goat; Abcam, ab36001, 1:400), and anti-TUJ1 (chicken; GeneTex, GTX85469, 1:200). Samples were washed three times for 10□min in PBS + 0.5% Triton X-100, then incubated in secondary antibody solution (secondary antibodies diluted in 5% normal donkey serum in PBS + 0.5% Triton X-100) for 4□h at room temperature or overnight at 4□°C. The following secondary antibodies were used: Alexa Fluor 488 Donkey anti-Chicken IgY (Thermo Fisher Scientific, A78948, 1:400), Alexa Fluor 647 Donkey anti-Rabbit IgG (Thermo Fisher Scientific, A-31573, 1:400), and Alexa Fluor 594 Donkey anti-Goat IgG (Thermo Fisher Scientific, A-11058, 1:400). Secondary antibody–incubated samples were washed three to five times for 10□min each in PBS + 0.5% Triton X-100, then mounted on slides using VECTASHIELD Antifade Mounting Medium without DAPI. Imaging was performed on a Nikon Sora Spinning Disk microscope, and quantification was conducted using Fiji.

#### Whole knee joints

For light sheet imaging of whole-knee joints, isolated knee joints were processed as previously described^50^. The following primary antibodies were used: anti-PGP9.5 (rabbit; Sigma, SAB4503057, 1:400) and anti-STEM121 (mouse; Takara, Y40410, 1:200). The following secondary antibodies were used: Alexa Fluor 488 Donkey anti-Mouse IgG (Thermo Fisher Scientific, A-21202, 1:500) and Alexa Fluor 594 Donkey anti-Rabbit IgG (Thermo Fisher Scientific, A-21207, 1:500). Imaging was performed on a Zeiss Lightsheet 7 Microscope, and quantification was conducted using Fiji.

### Safranin O-Fast Green Stain for Cartilage

Immediately following euthanasia, knee joint tissues were harvested from mice and fixed in 4% paraformaldehyde (PFA) for 48 hours. After fixation, samples underwent decalcification in 0.5M ethylenediaminetetraacetic acid (EDTA, pH 7.4) at room temperature for two weeks, followed by dehydration. Specimens were then embedded in paraffin and sectioned sagittally at a thickness of 4 μm and subjected to Safranin O-Fast Green staining according to standard protocols^11^.

### Behavior tests

Mechanical hypersensitivity was evaluated by the von Frey assay as previously described^22^. Mice were acclimated for at least 1 h on a wire-mesh platform beneath an opaque enclosure prior to testing. A calibrated 0.4 g von Frey filament (BIOSEB) was applied perpendicularly to the plantar surface of each hind paw and held for up to 3 s or until a brisk withdrawal response. Ten stimuli were delivered to each paw at 2 s intervals. The number of withdrawal responses per series was recorded, and three independent series were performed per paw. The mean withdrawal frequency across series was used for subsequent statistical analyses.

Knee joint mechanical hyperalgesia was measured using a SMALGO algometer (BIOSEB) as described^51^. Under gentle restraint, a 5 mm diameter probe was positioned against the distal femur adjacent to the knee joint. Force was increased at 50 g s⁻¹ from 0 g until an audible vocalization indicated withdrawal. Three trials were conducted per animal with ≥ 15 min rest intervals to prevent sensitization. The pressure at vocalization was recorded for each trial, and the average threshold was calculated. A 500 g cut-off was imposed to avoid tissue damage.

### microCT scanning and analysis

Knee joints were harvested and fixed in 10% neutral-buffered formalin for 24 h at room temperature. Following fixation, samples were washed in phosphate-buffered saline and stored in 70% ethanol until imaging. Fixed joints were scanned on a SkyScan 1275 high-resolution µCT system (Bruker) as previously described^51^. Acquisition parameters were set to 65 kV and 153 µA, yielding an isotropic voxel size of 6 µm. Raw projection images were reconstructed in NRecon v1.6 (Bruker) with beam-hardening correction and ring-artifact reduction. Three-dimensional segmentation and quantitative morphometry of the proximal tibial subchondral bone were performed in CTAn v1.9, and volumetric renderings were generated in CTVol v2.0 (Bruker). Representative sagittal sections are presented.

### Data Independent Acquisition (DIA) based Quantitative LC-MS/MS Proteomics

#### Sample Preparation

Protein extraction and digestion were performed based on an optimized protocol^52^. For the bone samples, approximately 50mg of bone joints were lysed in a denaturing buffer consisting of 8M urea, 50 mM HEPES (pH8.5) and 0.5% sodium deoxycholate at 4°C using a bullet blender. Protein concentration was determined using the BCA assay (Thermofisher Scientific). The lysates were digested with Lys-C (Wako, 1:100, w/w) for 3 h, followed by trypsin digestion (Promega, 1:50, w/w) overnight at room temperature. Prior to trypsin digestion, the solution was diluted 4 folds with 50 mM HEPES solution at pH 8.5. Reduction and alkylation were performed before acidifying the samples with 1% trifluoroacetic acid (TFA). Peptides were then desalted on C18 StageTip (Thermofisher Scientific), dried and stored at −80 ^0^C before MS analysis.

For human synovial fluid analysis, *CD200^high^* hPSC-NNs and *CD200^low^* hPSC-SNs were incubated overnight in OA synovial fluid (Biotrend, CDD-H-5000-OA) obtained from two independent donors distinct from those used in calcium imaging experiments. The supernatants were diluted with SDS buffer (2% SDS, 5□mM DTT, 50□mM HEPES at pH 8.5, and protease inhibitors) and then boiled at 95□°C for 10□minutes. The samples were then subjected to short gel SDS-PAGE and processed for in-gel digestion following an established protocol^52^. Briefly, protein bands were excised from Coomassie-stained gels, cut into ∼1□mm³ pieces, reduced with 5□mM DTT, alkylated with 10□mM IAA, and digested overnight at 37□°C. Following extraction, peptides were dried under vacuum and stored until further analysis.

#### LC-MS/MS Analysis

Proteomics analysis was performed using an Orbitrap Exploris 480 mass spectrometer (Thermo Scientific) coupled to an Ultimate 3000 (Thermo Scientific) HPLC operating in data-independent acquisition mode (DIA).

For mouse bone samples, the peptides were separated on a C18 column (20 cm × 75 μm, 1.7 μm particle size, CoANN technology) and eluted with a 13–48% buffer B gradient over 60 minutes (buffer A: 0.2% formic acid, 3% DMSO in water; buffer B: 0.2% formic acid, 3% DMSO in water in 67% acetonitrile) at a flow rate of 0.25□μL/min. The MS1 scan was acquired with a 60 K resolution within 500–1100 of m/z range, and the automatic gain control (AGC) value was set to 300% with 25 ms of maximum injection time (MIT). The MS2 scans were acquired with a 30 K resolution after fragmenting the precursor ions, which were obtained from 30 m/z isolation windows with high-energy collisional dissociation (HCD) collision energy as 30%.

For human synovial fluid samples, the peptides were separated on a C18 column (50 cm × 75 μm, 1.7 μm particle size, CoANN technology) and eluted with a 14–38% buffer B gradient over 60 minutes. The MS1 scan was acquired with a 60 K resolution within 350–1000 of m/z range, and the AGC value was set to 300% with 25 ms of MIT. The MS2 scans were acquired with a 30 K resolution after fragmenting the precursor ions, which are obtained from 30 isolation windows with HCD collision energy as 29%.

#### Database Search and Bioinformatic Analysis

The database searches were performed using DIA-NN (version 1.8) against an in-silico library that were generated from human (proteome ID: UP000005640) and mouse (proteome ID: UP000000589) protein sequences downloaded from the UniProt database^53^. Search parameters included: enzyme specificity set to Trypsin/P with up to 2 missed cleavages; carbamidomethylation of cysteines (+57.02146) as a fixed modification; methionine oxidation (+15.99491) as a variable modification (maximum of 2 variable modifications per peptide); precursor charge range of 1–4; precursor m/z range of 300–1,800; fragment m/z range of 200–1,800; and a precursor FDR threshold of 1%. Match-between-runs (MBR) was enabled. To eliminate potential cross-species contamination in mouse bone data, precursors mapping to both human and mouse proteins were excluded, and only mouse-specific precursors were retained for quantification. Quantitative analysis was performed following the same approach as described in our previous publication^54^. Briefly, for mouse datasets, the search results were further filtered with precursor q value of <0.01, and protein group q value of <0.01. For human datasets, the thresholds were set at precursor q value <0.05 and protein group q value <0.05. Protein group quantification was performed using the MaxLFQ algorithm implemented in the ‘iq’ R package^55^.

Differentially abundant proteins (DAPs) were identified using a custom R/Shiny application^56^. Missing values were imputed only when all the data in one group were completely missing while the other group had more than one observed value. In these scenarios, the missing values were imputed with the top n minimum values sample wisely in the group. The number of missing values imputed was equal to the maximum number of values in the other groups that had at least 2 values. Linear models for microarray data (Limma) were integrated into R/Shiny application to facilitate the identification of differentially abundant proteins (DAPs) in the proteomics dataset^57^. For mouse bone samples, differentially expressed proteins (DEPs) were identified using cutoffs |log2 fold change| ≥ 0.5 and P = 0.05. For human synovial fluids, DEPs were identified using cutoffs |log_2_ fold change| ≥ 0.2 and FDR < 0.05 to detect low quantities of sequestered or secreted proteins.

### Image analysis

All image analyses were performed using Fiji (v2.14.0). For quantification of ATF3, CGRP, STEM121, TNFα, COX2, CD206, CD86, and MMP13, one image was acquired per section per animal. Counts were performed independently by two individuals blinded to treatment conditions, and results were reported as the average of the two measurements. Quantification of total fiber length in knee joint sections was carried out using NeuronJ (v1.4.3). For whole knee joints imaged by light sheet microscopy, images were converted to 2D projections using maximum intensity projection. Binary thresholds were set uniformly across all images, and total nerve fiber length was quantified by applying skeletonization to the resulting binary images. Cartilage histopathology was scored based on prior grading the staging standards by two individuals blinded to treatment conditions^52^, and results were reported as the average of the two measurements.

### Statistical analysis

Statistical analyses were performed using GraphPad Prism v10.0.2. Comparisons were conducted using two-tailed unpaired Student’s t-tests, one-way ANOVA with Tukey’s multiple comparisons test, two-way ANOVA with Tukey’s multiple comparisons test, or two-way ANOVA with Sidak’s multiple comparisons test, unless otherwise indicated. Bar graphs depict mean□±□s.e.m. with individual data points overlaid; floating bars depict mean□±□range with individual data points overlaid.

### Data availability

All data is available upon request from the corresponding author. scRNA-seq and bulk RNA sequencing normalized counts, and proteomics data will be uploaded to NCBI upon publication.

## ACKNOWLEDGEMENTS

We thank Michael Caterina for his thoughtful input on the manuscript; Hojae Lee for early guidance in the maintenance and differentiation of hPSC lines; and Linda Orzolek, Tyler Creamer, Russell Hughes, and Jong-Seok Lee from the JHMI Single Cell and Transcriptomics Core for their support with sequencing. This work was supported by the Howard Hughes Medical Institute (to X.D.). X.D. and G.L. are supported by MSCRF grants 2022-MSCRFD-5872 and 2024-MSCRFV-6366. G.L. is also supported by NIH grant R21AI183368 and Blaustein Endowment for Pain Research and Education. X.D. is also supported by NIH grant R37NS054791. J.P. is supported by NIH grant RF1AG064909.

## AUTHOR CONTRIBUTIONS

Z.W., G.L., and X.D. conceptualized the study. Z.W. performed gene editing, differentiation and characterization of hPSC-SNs, calcium imaging and analysis, single-cell and bulk RNA-seq analysis, proteomics analysis, and tissue immunohistochemistry. W.Z. conducted animal surgeries, behavioral assays, and tissue immunohistochemistry. J.W. and Z.W. performed LC-MS/MS proteomics. X.C. reviewed data and provided input on animal studies. J.P. reviewed data and provided input on proteomics analysis. Z.W., G.L., and X.D. wrote the manuscript with input from all authors.

---

**Figure S1.**
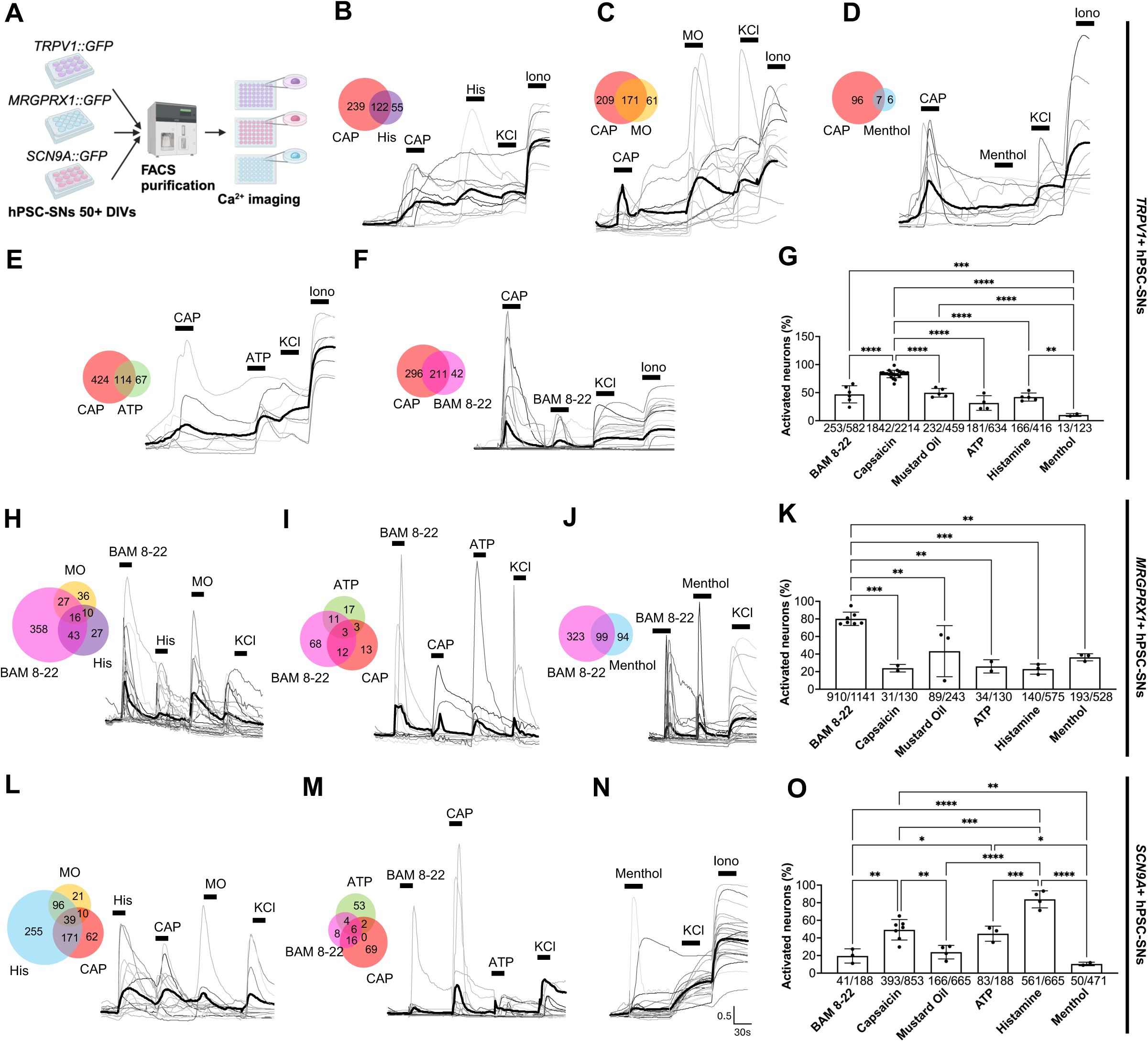
*SCN9A+, TRPV1+* nociceptors and *MRGPRX1+* pruriceptors are polymodal and respond to canonical noxious or pruritogenic stimuli, related to Figure 1. **(A)** Workflow of FACS purification of *MRGPRX1+, SCN9A+ and TRPV1+* hPSC-SNs followed by Ca^2+^ imaging. **(B-G)** Representative Ca^2+^ transients of FACS-purified *TRPV1*+ hPSC-SNs in response to capsaicin (10μM; **B-F**), histamine (50μM; **B**), mustard oil (100μM; **C**), menthol (100μM; **D**), ATP (50μM; **E**), and BAM8-22 (10μM; **F**); bold traces show the mean response. Venn diagrams show the overlap in stimulus responsiveness. Quantification of proportion of *TRPV1*+ neurons activated by each stimuli **(G)** (n ≥ 3 independent differentiations; one-way ANOVA with Tukey’s multiple comparisons test; ** P < 0.01, *** P < 0.001, **** P < 0.0001; mean ± s.e.m. with individual values overlaid). **(H-K)** Representative Ca^2+^ transients of FACS-purified *MRGPRX1*+ hPSC-SNs in response to capsaicin (10μM; **I**), histamine (50μM; **H**), mustard oil (100μM; **H**), menthol (100μM; **J**), ATP (50μM; **I**), and BAM8-22 (10μM; **H-J**); bold traces show the mean response. Venn diagrams show the overlap in stimulus responsiveness. Quantification of proportion of *MRGPRX1*+ neurons activated by each stimuli **(K)** (n ≥ 3 independent differentiations; one-way ANOVA with Tukey’s multiple comparisons test; ** P < 0.01, *** P < 0.001; mean ± s.e.m. with individual values overlaid). **(L-O)** Representative Ca^2+^ transients of FACS-purified *SCN9A*+ hPSC-SNs in response to capsaicin (10μM; **L,M**), histamine (50μM; **L**), mustard oil (100μM; **L**), menthol (100μM; **N**), ATP (50μM; **M**), and BAM8-22 (10μM; **M**); bold traces show the mean response. Venn diagrams show the overlap in stimulus responsiveness. Quantification of proportion of *SCN9A*+ neurons activated by each stimuli **(O)** (n ≥ 3 independent differentiations; one-way ANOVA with Tukey’s multiple comparisons test; * P < 0.05, ** P < 0.01, *** P < 0.001, **** P < 0.0001; mean ± s.e.m. with individual values overlaid).

**Figure S2.**
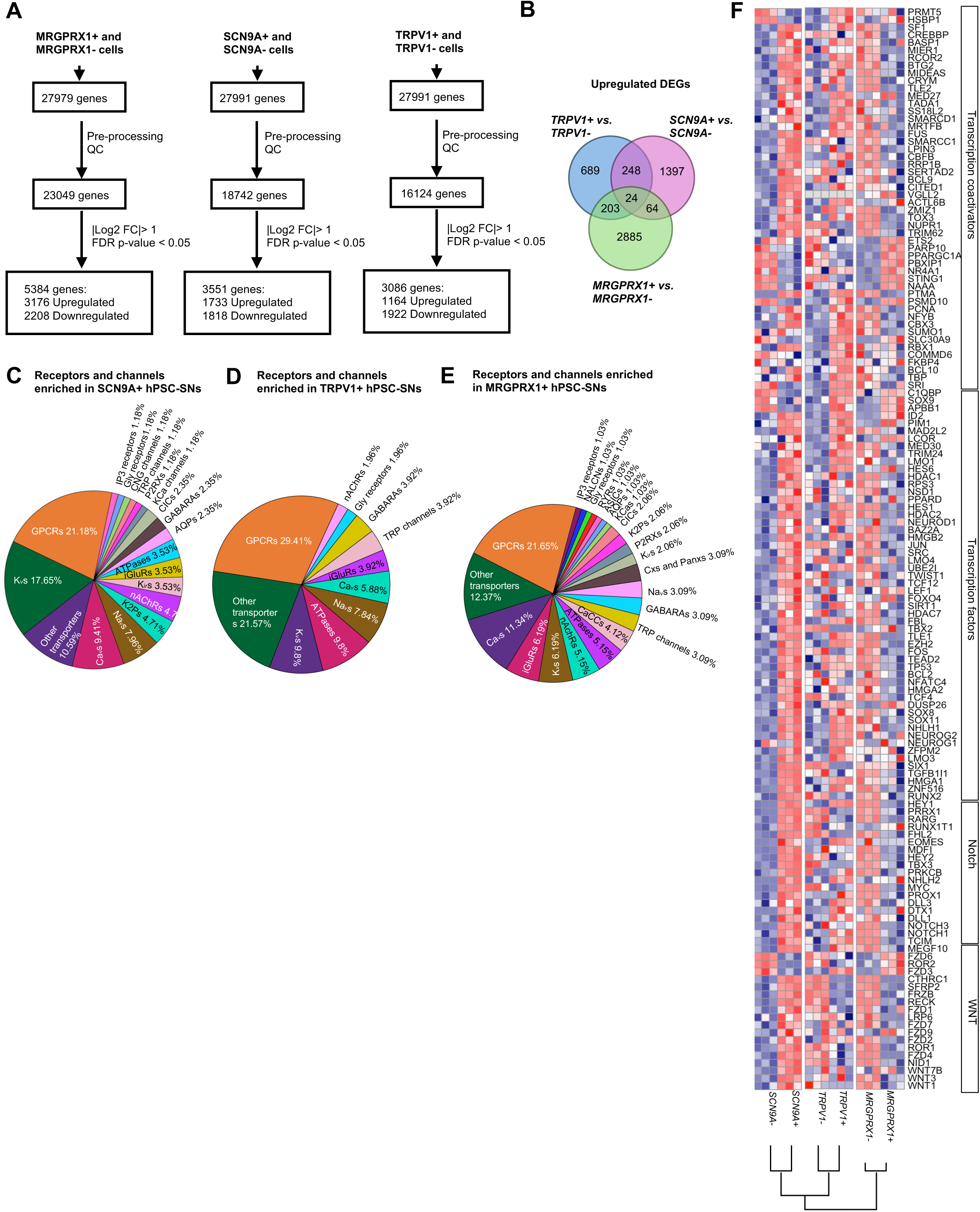
Transcriptomic characterization of *SCN9A+, TRPV1+* nociceptors and *MRGPRX1+* pruriceptors, related to Figure 2. **(A)** Workflow of purifying GFP+ and GFP-cells from hPSC-SN cultures of *MRGPRX1::GFP, SCN9A::GFP*, and *TRPV1::GFP* reporter lines (n = 5 independent differentiations; n = 3 technical replicates; log_₂_FC cutoff = 1; P cutoff = 0.05). **(B)** Venn diagram shows significantly enriched genes in *MRGPRX1+, SCN9A+ and TRPV1+* hPSC-SNs compared to their negative counterparts, highlighting shared and lineage-specific genes. **(C-E)** Pie charts categorizing enriched receptors and ion channels enriched in *SCN9A*+ **(C)**, *TRPV1*+ **(D)** and *MRGPRX1*+ **(E)** hPSC-SNs. **(F)** GSEA on differentially expressed transcription factors and key pathway proteins between GFP+ and GFP-populations in each reporter line using MSigDB gene sets; hierarchical clustering of pairwise lineage comparisons is shown as a dendrogram.

**Figure S3.**
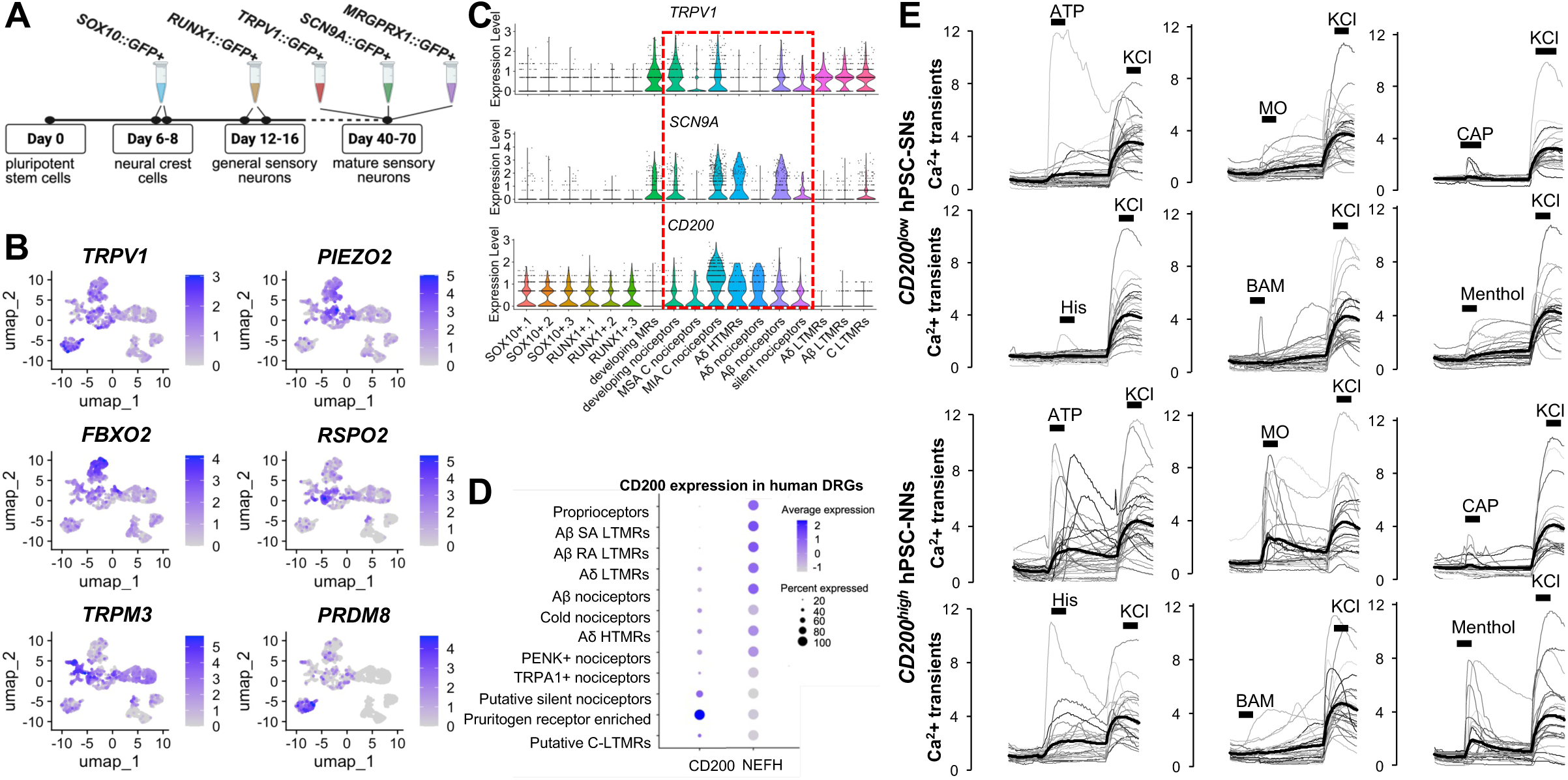
Transcriptomic and functional analyses reveal specific enrichment of *CD200* in nociceptors, related to Figure 3. **(A)** Workflow for constructing a transcriptomic continuum of nociceptor development from FACS-purified *SOX10::GFP*+ neural crest cells, *RUNX1::GFP*+ sensory progenitor cells, *MRGPRX1::GFP*+, *SCN9A::GFP*+, and *TRPV1::GFP*+ hPSC-SNs by scRNA-seq (≈ 1,000 cells per lineage; n = 3 independent differentiations). **(B)** Feature plots on the UMAP embedding showing expression of additional marker genes used to assign each cluster identity. **(C)** Violin plots showing the co-enrichment of *TRPV1*, *SCN9A* and *CD200* in nociceptor clusters. **(D)** Dot plot showing *CD200* enrichment in small-fiber nociceptors form a published human DRG neuron single-cell dataset^7^. **(E)** Representative Ca^2+^ transients demonstrating heightened responses of *CD200^high^* hPSC-NNs versus *CD200^low^* hPSC-SNs to ATP (50μM), mustard oil (100μM), capsaicin (10μM), and histamine (50μM); bold traces show the mean response; n ≥ 3 independent differentiations. The response rate to BAM8-22 (10 μM) and menthol (100 μM) did not show significant differences.

**Figure S4.**
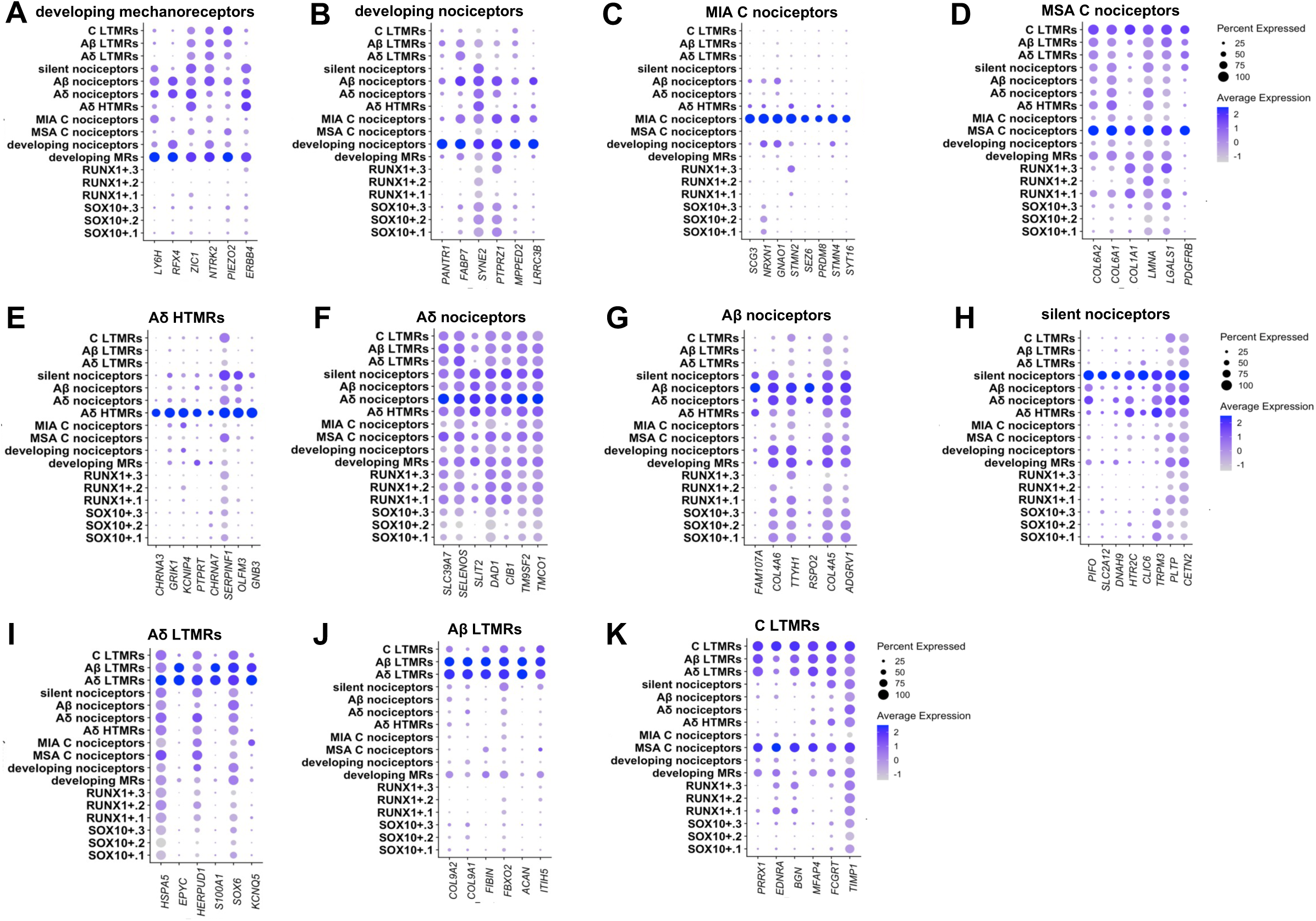
Marker gene expression across hPSC-SNs clusters, related to Figure 2. **(A-K)** Dot plots of the top marker genes for each identified neuronal subpopulation, showing their expression across all clusters; dot size reflects the fraction of cells within each cluster expressing the gene, and dot color indicates the mean scaled expression level.

**Figure S5.**
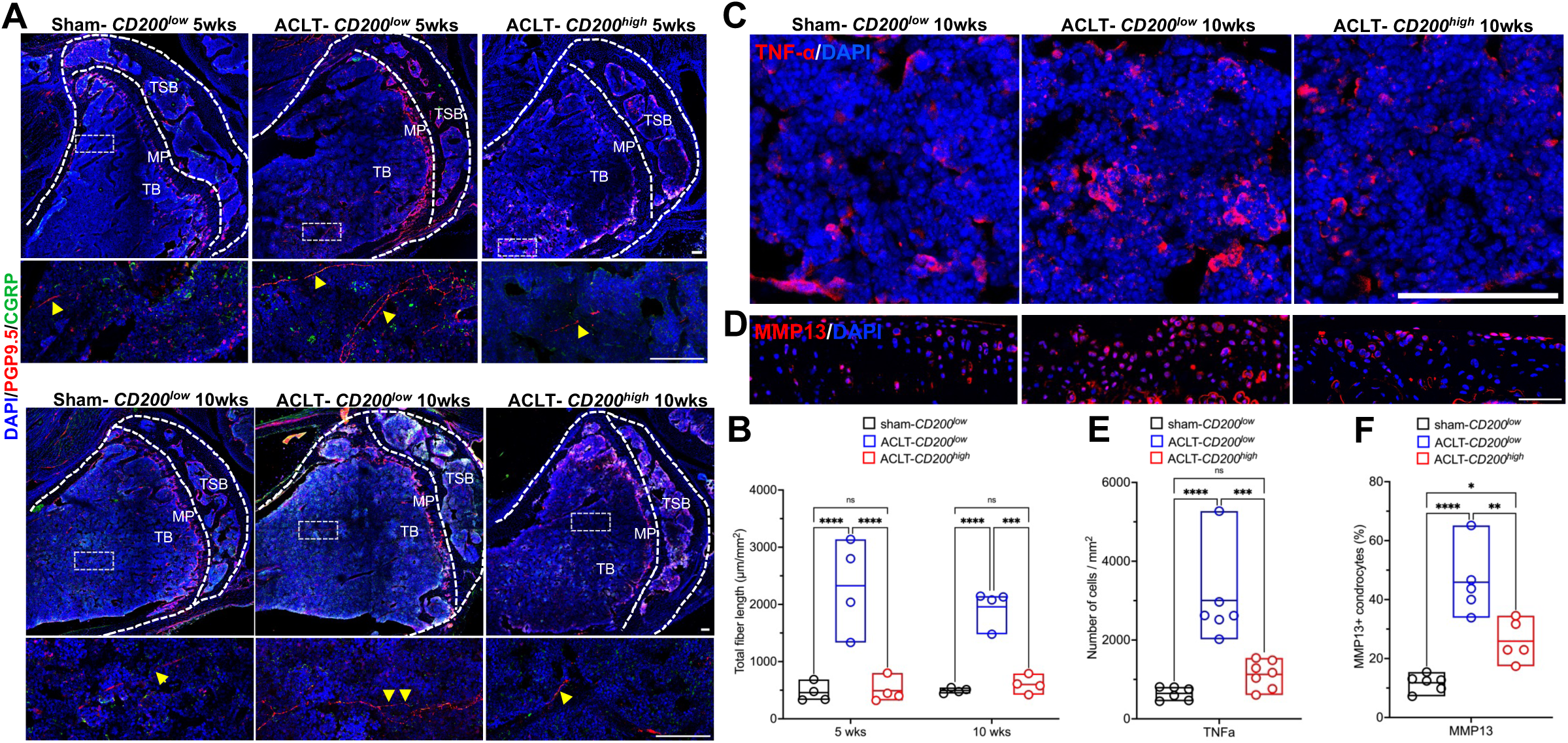
*CD200^high^*hPSC-NNs modulate the neuro-immune axis in ACLT-*CD200^high^* versus ACLT-*CD200^low^* animals, related to Figure 4. **(A)** PGP9.5 (red) and CGRP (green) immunostaining in knee sections from sham-*CD200^low^*, ACLT-*CD200^low^* and ACLT-*CD200^high^* animals 5 (top) and 10 (bottom) weeks post-injection; scale bars, 100µm. TSP, tibial subchondral plate; MP, metaphysis; TB, tibial bone. **(B)** Quantification of total PGP9.5+ nerve fiber length 5 and 10 weeks post-injection (n ≥ 4 animals; two-way ANOVA with Tukey’s test; *** P < 0.001, **** P < 0.0001; mean ± range with individual values overlaid). **(C,E)** TNF-α (red) immunostaining in sham-*CD200^low^*, ACLT-*CD200^low^*and ACLT-*CD200^high^* joints 10 weeks post-injection; scale bar, 100µm. Quantification of TNF-α expression **(E)** (n ≥ 6 animals; one-way ANOVA with Tukey’s test; *** P < 0.001, **** P < 0.0001; mean ± range with individual values overlaid). **(D,F)** MMP13 (red) immunostaining in sham-*CD200^low^*, ACLT-*CD200^low^* and ACLT-*CD200^high^* joints 10 weeks post-injection; scale bar, 100µm. Quantification of MMP13 expression **(F)** (n ≥ 5 animals; one-way ANOVA with Tukey’s test; * P < 0.05, ** P < 0.01, **** P < 0.0001; mean ± range with individual values overlaid).

**Figure S6.**
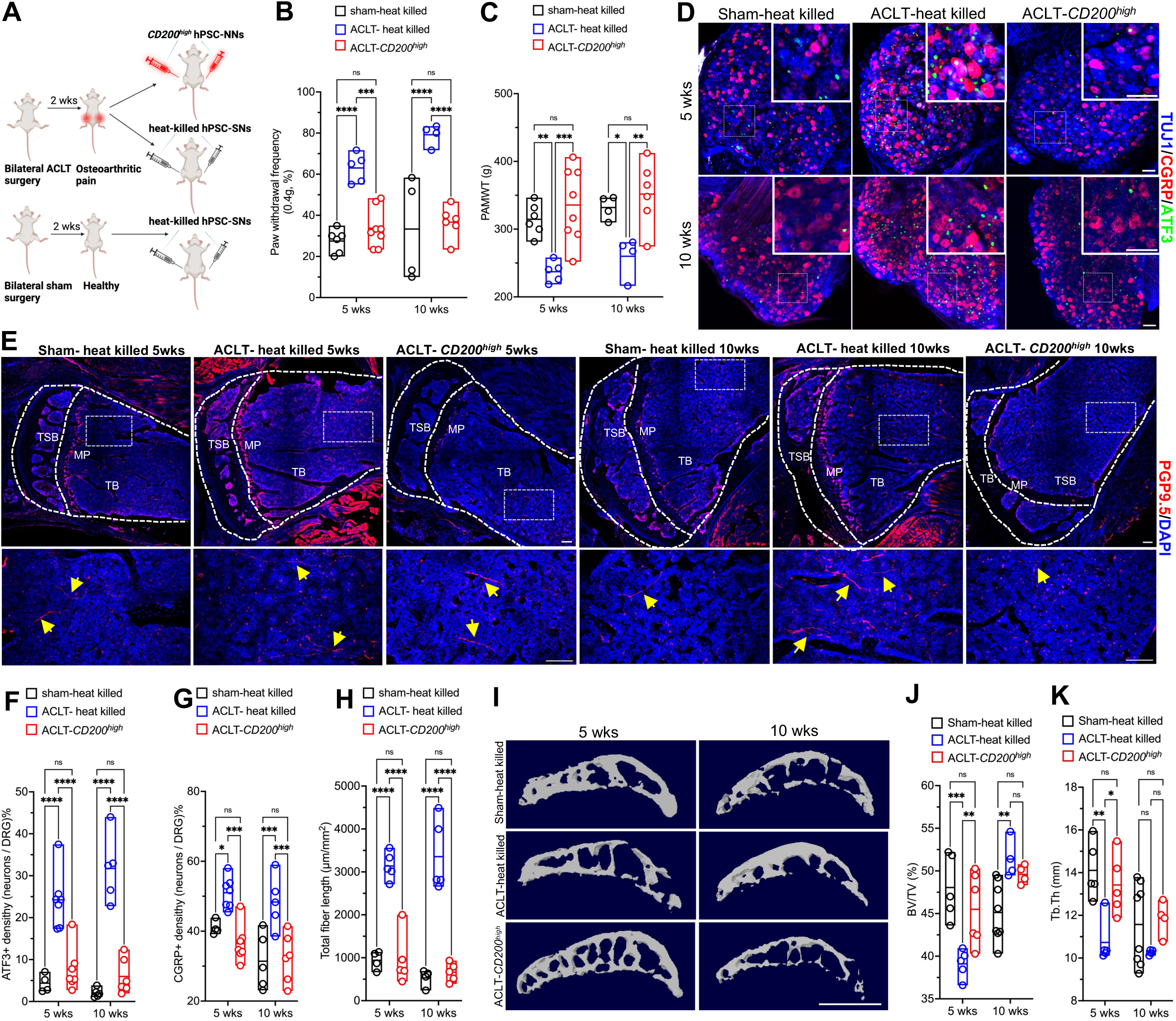
*CD200^high^* hPSC-NNs reduce pain and improve subchondral microarchitecture in ACLT-*CD200^high^* versus ACLT-heat killed animals, related to Figure 4. **(A)** Experimental timeline of bilateral intra-articular and tibia tuberosity injections of *CD200^high^* hPSC-NNs or heat-killed hPSC-SNs in NIH-III mice two weeks after sham or ACLT surgery. **(B-C)** Mechanical hypersensitivity assessed by Von Frey **(B)** and PAMWT **(C)** in sham-heat killed, ACLT-heat killed and ACLT-*CD200^high^* animals 5 and 10 weeks post-injection (n ≥ 4; two-way ANOVA with Tukey’s multiple comparisons test; * P < 0.05, ** P < 0.01, *** P < 0.001, **** P < 0.0001; mean ± range with individual values overlaid). **(D,F,G)** L3–L4 whole DRGs immunostained for ATF3 (green) and CGRP (red) at 5 and 10 weeks; scale bars, 100µm. Quantification of ATF3+ **(F)** and CGRP+ **(G)** neurons in whole DRGs (n ≥ 5; two-way ANOVA with Tukey’s test; *** P < 0.001, **** P < 0.0001; mean ± range with individual values overlaid). **(E)** PGP9.5 immunostaining (red, arrows) in knee sections from sham-heat killed, ACLT-heat killed and ACLT-*CD200^high^*animals 5 (left) and 10 (right) weeks post-injection; scale bars, 100µm. TSP, tibial subchondral plate; MP, metaphysis; TB, tibial bone. **(H)** Quantification of total PGP9.5+ nerve fiber length 5 and 10 weeks post-injection (n ≥ 4 animals; two-way ANOVA with Tukey’s test; **** P < 0.0001; mean ± range with individual values overlaid). **(I-K)** Representative microCT of tibia subchondral bone **(I)** and quantification of trabecular bone volume fraction (BV/TV; **J**) and trabecular thickness (Tb.Th; **K**) (n ≥ 5; two-way ANOVA with Tukey’s test; * P < 0.05, ** P < 0.01, *** P < 0.001; mean ± range with individual values overlaid).

## REFERENCE

1. Pezet, S. & McMahon, S. B. Neurotrophins: mediators and modulators of pain. Annu. Rev. Neurosci. 29507–538. 10.1146/annurev.neuro.29.051605.112929 (2006)

2. Chen, D. et al. Osteoarthritis: toward a comprehensive understanding of pathological mechanism. Bone Res 516044. 10.1038/boneres.2016.44 (2017)

3. da Costa, B. R. et al. Effectiveness and safety of non-steroidal anti-inflammatory drugs and opioid treatment for knee and hip osteoarthritis: network meta-analysis. BMJ (Clinical research ed.) 375n2321. 10.1136/bmj.n2321 (2021)

4. Sanga, P. et al. Efficacy, safety, and tolerability of fulranumab, an anti-nerve growth factor antibody, in the treatment of patients with moderate to severe osteoarthritis pain. Pain 154(10), 1910–1919. 10.1016/j.pain.2013.05.051 (2013)

5. Tabar, V. & Studer, L. Pluripotent stem cells in regenerative medicine: challenges and recent progress. Nat. Rev. Genet. 15(2), 82–92. 10.1038/nrg3563 (2014)

6. Chambers, S. M., et al. Highly efficient neural conversion of human ES and iPS cells by dual inhibition of SMAD signaling. Nat. Biotechnol. 27(3), 275–280. 10.1038/nbt.1529 (2009)

7. Nguyen, M. Q., von Buchholtz, L. J., Reker, A. N., Ryba, N. J. & Davidson, S. Single-nucleus transcriptomic analysis of human dorsal root ganglion neurons. Elife 10e71752. 10.7554/eLife.71752 (2021)

8. Tavares-Ferreira, D. et al. Spatial transcriptomics of dorsal root ganglia identifies molecular signatures of human nociceptors. Sci. Transl. Med. 14(632), eabj8186. 10.1126/scitranslmed.abj8186 (2022)

9. Li, Z. et al. Targeting human Mas-related G protein-coupled receptor X1 to inhibit persistent pain. Proc. Natl. Acad. Sci. U. S. A 114(10), E1996–E2005. 10.1073/pnas.1615255114 (2017)

10. Caterina, M. J. et al. The capsaicin receptor: a heat-activated ion channel in the pain pathway. Nature 389(6653), 816–824. 10.1038/39807 (1997)

11. Facer, P. et al. Differential expression of the capsaicin receptor TRPV1 and related novel receptors TRPV3, TRPV4 and TRPM8 in normal human tissues and changes in traumatic and diabetic neuropathy. BMC Neurol. 711. 10.1186/1471-2377-7-11 (2007)

12. Schwartzentruber, J. et al. Molecular and functional variation in iPSC-derived sensory neurons. Nat. Genet. 50(1), 54–61. 10.1038/s41588-017-0005-8 (2018)

13. Hao, Y. et al. Integrated analysis of multimodal single-cell data. Cell 184(13), 3573–3587.e29. 10.1016/j.cell.2021.04.048 (2021)

14. Yu, H. et al. Single-Soma Deep RNA sequencing of Human DRG Neurons Reveals Novel Molecular and Cellular Mechanisms Underlying Somatosensation. bioRxiv : the preprint server for biology 2023.03.17.533207 [pii]. 10.1101/2023.03.17.533207 (2023)

15. Cui, M. & Nicol, G. D. Cyclic AMP mediates the prostaglandin E2-induced potentiation of bradykinin excitation in rat sensory neurons. Neuroscience 66(2), 459–466. 10.1016/0306-4522(94)00567-o (1995)

16. Wirotanseng, L. N., Kuner, R. & Tappe-Theodor, A. Gq rather than G11 preferentially mediates nociceptor sensitization. Mol. Pain 954. 10.1186/1744-8069-9-54 (2013)

17. Qian, J. J., Xu, Q., Xu, W. M., Cai, R. & Huang, G. C. Expression of VEGF-A Signaling Pathway in Cartilage of ACLT-induced Osteoarthritis Mouse Model. J. Orthop. Surg. Res. 16(1), 379. 10.1186/s13018-021-02528-w (2021)

18. Kelly, S. et al. Transplanted human fetal neural stem cells survive, migrate, and differentiate in ischemic rat cerebral cortex. Proc. Natl. Acad. Sci. U. S. A 101(32), 11839–11844. 10.1073/pnas.0404474101 (2004)

19. Bráz, J. M. & Basbaum, A. I. Differential ATF3 expression in dorsal root ganglion neurons reveals the profile of primary afferents engaged by diverse noxious chemical stimuli. Pain 150(2), 290–301. 10.1016/j.pain.2010.05.005 (2010)

20. Seijffers, R., Mills, C. D. & Woolf, C. J. ATF3 increases the intrinsic growth state of DRG neurons to enhance peripheral nerve regeneration. J. Neurosci. 27(30), 7911–7920. 10.1523/JNEUROSCI.5313-06.2007 (2007)

21. Staton, P. C., Wilson, A. W., Bountra, C., Chessell, I. P. & Day, N. C. Changes in dorsal root ganglion CGRP expression in a chronic inflammatory model of the rat knee joint: differential modulation by rofecoxib and paracetamol. European journal of pain (London, England) 11(3), 283–289. 10.1016/j.ejpain.2006.03.006 (2007)

22. Zhang, W. et al. RANK(+)TLR2(+) myeloid subpopulation converts autoimmune to joint destruction in rheumatoid arthritis. Elife 12e85553. 10.7554/eLife.85553 (2023)

23. Zhu, S. et al. Subchondral bone osteoclasts induce sensory innervation and osteoarthritis pain. J. Clin. Invest. 129(3), 1076–1093. 10.1172/JCI121561 (2019)

24. Loeser, R. F., Goldring, S. R., Scanzello, C. R. & Goldring, M. B. Osteoarthritis: a disease of the joint as an organ. Arthritis and rheumatism 64(6), 1697–1707. 10.1002/art.34453 (2012)

25. Pecchi, E. et al. Induction of nerve growth factor expression and release by mechanical and inflammatory stimuli in chondrocytes: possible involvement in osteoarthritis pain. Arthritis Res. Ther. 16(1), R16. 10.1186/ar4443 (2014)

26. Deng, H., Li, J., Shah, A. A., Ge, L. & Ouyang, W. Comprehensive in-silico analysis of deleterious SNPs in APOC2 and APOA5 and their differential expression in cancer and cardiovascular diseases conditions. Genomics 115(2), 110567. 10.1016/j.ygeno.2023.110567 (2023)

27. Zewinger, S. et al. Apolipoprotein C3 induces inflammation and organ damage by alternative inflammasome activation. Nat. Immunol. 21(1), 30–41. 10.1038/s41590-019-0548-1 (2020)

28 . Appleton, C. T., Usmani, S. E., Mort, J. S. & Beier, F. Rho/ROCK and MEK/ERK activation by transforming growth factor-alpha induces articular cartilage degradation. Lab. Invest. 90(1), 20–30. 10.1038/labinvest.2009.111 (2010)

29 . Ashraf, S., et al. Increased vascular penetration and nerve growth in the meniscus: a potential source of pain in osteoarthritis. Ann. Rheum. Dis. 70(3), 523–529. 10.1136/ard.2010.137844 (2011)

30. Wang, J. et al. Altered expression of microRNA-98 in IL-1β-induced cartilage degradation and its role in chondrocyte apoptosis. Mol. Med. Report. 16(3), 3208–3216. 10.3892/mmr.2017.7028 (2017)

31. Maumus, M. et al. Thrombospondin-1 Partly Mediates the Cartilage Protective Effect of Adipose-Derived Mesenchymal Stem Cells in Osteoarthritis. Front. Immunol. 81638. 10.3389/fimmu.2017.01638 (2017)

32. Meng, C. et al. Therapeutic potential of CHI3L1 in osteoarthritis: Inhibition of cartilage matrix degradation and inflammation through TLR4-MAPK-STAT1 pathway. Int. Immunopharmacol. 156114684. 10.1016/j.intimp.2025.114684 (2025)

33. Luo, S., Li, W., Wu, W. & Shi, Q. Elevated expression of MMP8 and MMP9 contributes to diabetic osteoarthritis progression in a rat model. J. Orthop. Surg. Res. 16(1), 64. 10.1186/s13018-021-02208-9 (2021)

34. Ruettger, A., Schueler, S., Mollenhauer, J. A. & Wiederanders, B. Cathepsins B, K, and L are regulated by a defined collagen type II peptide via activation of classical protein kinase C and p38 MAP kinase in articular chondrocytes. JOURNAL OF BIOLOGICAL CHEMISTRY 283(2), 1043–1051. 10.1074/jbc.M704915200 (2008)

35. Shimoide, T. et al. Roles of Olfactomedin 1 in Muscle and Bone Alterations Induced by Gravity Change in Mice. Calcif. TIssue Int. 107(2), 180–190. 10.1007/s00223-020-00710-6 (2020)

36. Heinegård, D. & Saxne, T. The role of the cartilage matrix in osteoarthritis. Nat Rev Rheumatol 7(1), 50–56. 10.1038/nrrheum.2010.198 (2011)

37. Ertürk, C., Altay, M. A., Selek, S. & Koçyiğit, A. Paraoxonase-1 activity and oxidative status in patients with knee osteoarthritis and their relationship with radiological and clinical parameters. Scand. J. Clin. Lab. Invest. 72(5), 433–439. 10.3109/00365513.2012.687116 (2012)

38. Kirkeby, A., Main, H. & Carpenter, M. Pluripotent stem-cell-derived therapies in clinical trial: A 2025 update. Cell Stem Cell 32(1), 10–37. 10.1016/j.stem.2024.12.005 (2025)

39. Ansari, A. M. et al. Cellular GFP Toxicity and Immunogenicity: Potential Confounders in in Vivo Cell Tracking Experiments. Stem Cell Rev Rep 12(5), 553–559. 10.1007/s12015-016-9670-8 (2016)

40. Chao, M. V. Neurotrophins and their receptors: a convergence point for many signalling pathways. Nat. Rev. Neurosci. 4(4), 299–309. 10.1038/nrn1078 (2003)

41. Crane, J. et al. Slit3 by PTH-Induced Osteoblast Secretion Repels Sensory Innervation in Spine Porous Endplates to Relieve Low Back Pain. Research square rs.3.rs-4823095 [pii]. 10.21203/rs.3.rs-4823095/v1 (2024)

42. Willis, C. M. et al. Harnessing the Neural Stem Cell Secretome for Regenerative Neuroimmunology. Front. Cell. Neurosci. 14590960. 10.3389/fncel.2020.590960 (2020)

43. Li, Y. et al. Rheumatoid arthritis synovial fluid induces JAK-dependent intracellular activation of human sensory neurons. JCI Insight 10(12), e186646 [pii]. 10.1172/jci.insight.186646 (2025)

44. Ng, Y. P., Cheung, Z. H. & Ip, N. Y. STAT3 as a downstream mediator of Trk signaling and functions. JOURNAL OF BIOLOGICAL CHEMISTRY 281(23), 15636–15644. 10.1074/jbc.M601863200 (2006)

45. Atri, D. S., et al. CRISPR-Cas9 Genome Editing of Primary Human Vascular Cells In Vitro. Current protocols 1(11), e291. 10.1002/cpz1.291 (2021)

46. Paquet, D. et al. Efficient introduction of specific homozygous and heterozygous mutations using CRISPR/Cas9. Nature 533(7601), 125–129. 10.1038/nature17664 (2016)

47. Björnson, Y. et al. The effect of histone deacetylase inhibitors on the efficiency of the CRISPR/Cas9 system. Biochem Biophys Rep 35101513. 10.1016/j.bbrep.2023.101513 (2023)

48. Umehara, Y. et al. Robust induction of neural crest cells to derive peripheral sensory neurons from human induced pluripotent stem cells. Sci. Rep. 10(1), 4360. 10.1038/s41598-020-60036-z (2020)

49. Wang, X. et al. Inhibition of Integrin αvβ6 Activation of TGF-β Attenuates Tendinopathy. Advanced science (Weinheim, Baden-Wurttemberg, Germany) 9(11), e2104469. 10.1002/advs.202104469 (2022)

50 . Ko, F. C., et al. Clearing-enabled light sheet microscopy as a novel method for three-dimensional mapping of the sensory innervation of the mouse knee. J. Orthop. Res. 43(3), 632–639. 10.1002/jor.26016 (2025)

51. Deng, R. et al. Periosteal CD68+ F4/80+ Macrophages Are Mechanosensitive for Cortical Bone Formation by Secretion and Activation of TGF-β1. Advanced science (Weinheim, Baden-Wurttemberg, Germany) 9(3), e2103343. 10.1002/advs.202103343 (2022)

52. Pagala, V. R. et al. Quantitative protein analysis by mass spectrometry. Methods in molecular biology (Clifton, N.J.) 1278281–305. 10.1007/978-1-4939-2425-7_17 (2015)

53. Demichev, V., Messner, C. B., Vernardis, S. I., Lilley, K. S. & Ralser, M. DIA-NN: neural networks and interference correction enable deep proteome coverage in high throughput. Nat. Methods 17(1), 41–44. 10.1038/s41592-019-0638-x (2020)

54. Wang, J. et al. Evaluation of Protein Identification and Quantification by the diaPASEF Method on timsTOF SCP. J. Am. Soc. Mass Spectrom. 35(6), 1253–1260. 10.1021/jasms.4c00067 (2024)

55. Pham, T. V., Henneman, A. A. & Jimenez, C. R. iq: an R package to estimate relative protein abundances from ion quantification in DIA-MS-based proteomics. Bioinformatics 36(8), 2611–2613. 10.1093/bioinformatics/btz961 (2020)

56. Kong, D. et al. A computational tool to infer enzyme activity using post-translational modification profiling data. Commun Biol 8(1), 103. 10.1038/s42003-025-07548-4 (2025)

57. Ritchie, M. E. et al. limma powers differential expression analyses for RNA-sequencing and microarray studies. Nucleic Acids Res. 43(7), e47. 10.1093/nar/gkv007 (2015)

